# Adipocyte *microRNA-802* promotes adipose tissue inflammation and insulin resistance by modulating macrophages in obesity

**DOI:** 10.1101/2024.05.13.593999

**Authors:** Yue Yang, Bin Huang, Yimeng Qin, Danwei Wang, Yinuo Jin, Linmin Su, Yi Pan, Yanfeng Zhang, Yumeng Shen, Wenjun Hu, Zhengyu Cao, Liang Jin, Fangfang Zhang

## Abstract

Adipose tissue inflammation is now considered to be a key process underlying metabolic diseases in obese individuals. However, it remains unclear how adipose inflammation is initiated and maintained or the mechanism by which inflammation develops. We found that *microRNA-802* (*miR-802*) expression in adipose tissue is progressively increased with the development of dietary obesity in obese mice and humans. The increasing trend of *miR-802* preceded the accumulation of macrophages. Adipose tissue-specific knockout of *miR-802* lowered macrophage infiltration and ameliorated systemic insulin resistance. Conversely, the specific overexpression of *miR-802* in adipose tissue aggravated adipose inflammation in mice fed a high-fat diet. Mechanistically, *miR-802* activates noncanonical and canonical NF-κB pathways by targeting its negative regulator, TRAF3. Next, NF-κB orchestrated the expression of chemokine and SREBP1, which translated into strong recruitment and M1-like polarization of macrophages. Our findings indicate that *miR-802* endows adipose tissue with the ability to recruit and polarize macrophages, which underscores *miR-802* as an innovative and attractive candidate for miRNA-based immune therapy for adipose inflammation.

## Introduction

Obesity is a very powerful health determinant or indicator that facilitates the development and progression of several metabolic diseases, including insulin resistance and type 2 diabetes(1, 2). Adipose tissue is a highly dynamic metabolic organ that plays a central role in the regulation of energy homeostasis and controls glucose metabolism and insulin sensitivity(3, 4). A hallmark of obesity is low-grade chronic inflammation in adipose tissue, marked by the accumulation of macrophages and other immune cells and by an increase in the levels of pro-inflammatory cytokines(5-7). Persistent adipose tissue inflammation is now considered to have a pivotal role in obesity-associated insulin resistance(8, 9). Resetting the immunological balance in obesity could represent an innovative approach for the management of insulin resistance and diabetes(10, 11). However, the early triggers and signals that sustain adipose tissue inflammation in obesity remain elusive, limiting our ability to effectively intervene this growing public health issue.

Macrophages accumulate in the adipose tissue of obese mice and humans, where they form crown-like structures surrounding dying or dead adipocytes and are key contributors to inflammation and obesity-induced insulin resistance(12, 13). The number of adipose tissue macrophages is tightly linked to the degree of insulin resistance and metabolic dysregulation(14, 15). Ablation of pro-inflammatory adipose tissue macrophages leads to a rapid improvement in insulin sensitivity and glucose tolerance, associated with marked decreases in local and systemic inflammation in obese mice(16, 17). Targeting the major inflammatory pathways is sufficient to counteract obesity-related systemic inflammation and insulin resistance(18, 19). However, the molecular links between lipid-overloaded adipocytes and inflammatory macrophages in obese adipose tissue remain elusive.

MicroRNAs (miRNAs) are small non-coding RNAs that post transcriptionally regulate gene expression by binding to specific regions of target genes to prevent translation or promote mRNA degradation(20). Emerging evidence suggests that miRNAs are key regulators in a variety of important metabolic organs and substantial contributors to the pathogenesis of complex diseases, including obesity-associated metabolic diseases(21, 22). In the adipose tissue, miRNAs have dramatic effects on regulating the pathways that control a range of processes including lipogenesis, inflammation, and insulin signaling(23, 24). Moreover, mice with alterations in the levels of miRNAs in adipocytes show significantly enhanced inflammation and insulin resistance after feeding with a high-fat diet (HFD), further confirming the contribution of miRNAs to obesity-induced phenotypes(25, 26). Therefore, adipose-derived miRNAs hold great promise for understanding adipose tissue dysfunction and the relationship between chronic inflammation and obesity and insulin resistance.

In this study, we demonstrated that *microRNA-802* (*miR-802*) promotes inter-cellular communication between lipid-overloaded adipocytes and macrophages, ultimately leading to adipose tissue inflammation and insulin resistance. Adipocyte *miR-802* levels are positively associated with obesity in mice and humans. Adipose tissue-specific overexpression of *miR-802* in mice fed an HFD exhibited increased severity of systemic insulin resistance compared with wild-type (WT) mice, which was accompanied by macrophage infiltration and a marked increase in adipose tissue inflammation. Adipose tissue-specific knockout of *miR-802* achieved the opposite result. Co-culture and other *in vitro* experiments revealed a vicious cycle of interactions between macrophages and adipocytes ectopically expressing *miR-802*. We established that *miR-802* expression is an inflammatory signal in adipocytes, and this effect occurs through sensitization of the NF-κB signaling pathway. Altogether, our data raise the possibility that manipulation of this microRNA action axis has therapeutic potential for treating adipose inflammation.

## Results

### *miR-802* elevation precedes macrophage accumulation

Consistent with previous studies from our and other laboratories(27, 28), adipose from obesity mice showed significantly higher *miR-802* expression than those from normal mice. To evaluate whether *miR-802* is involved in adipose inflammation and insulin resistance, we examined the expression profile of *miR-802*. we observed that *miR-802* progressively increased in adipose tissue from week 4 with the development of obesity in mouse models of genetic and dietary obesity (Figure 1A, B and Figure S1A, B). We next compared the expression of *miR-802* in different adipose depots and found that it was the highest in epididymal white adipose tissue (epiWAT) (Figure 1C). We further isolated mature adipose tissue and stromal vascular fraction (SVF) from epiWAT to examine the expression of *miR-802*. We found that *miR-802* expression was substantially higher in mature adipocytes than in SVF in both mice fed a normal chow diet (NCD) and those fed an HFD (Figure 1D). Through *in vitro* experiments, we found that *miR-802* was dramatically increased in the insulin resistance cell models (Figure 1E, F and Figure S1C, D). These findings suggest that upregulation of *miR-802* in adipocytes may be functionally involved in the pathogenesis of obesity-associated disorders.

**Figure 1.**
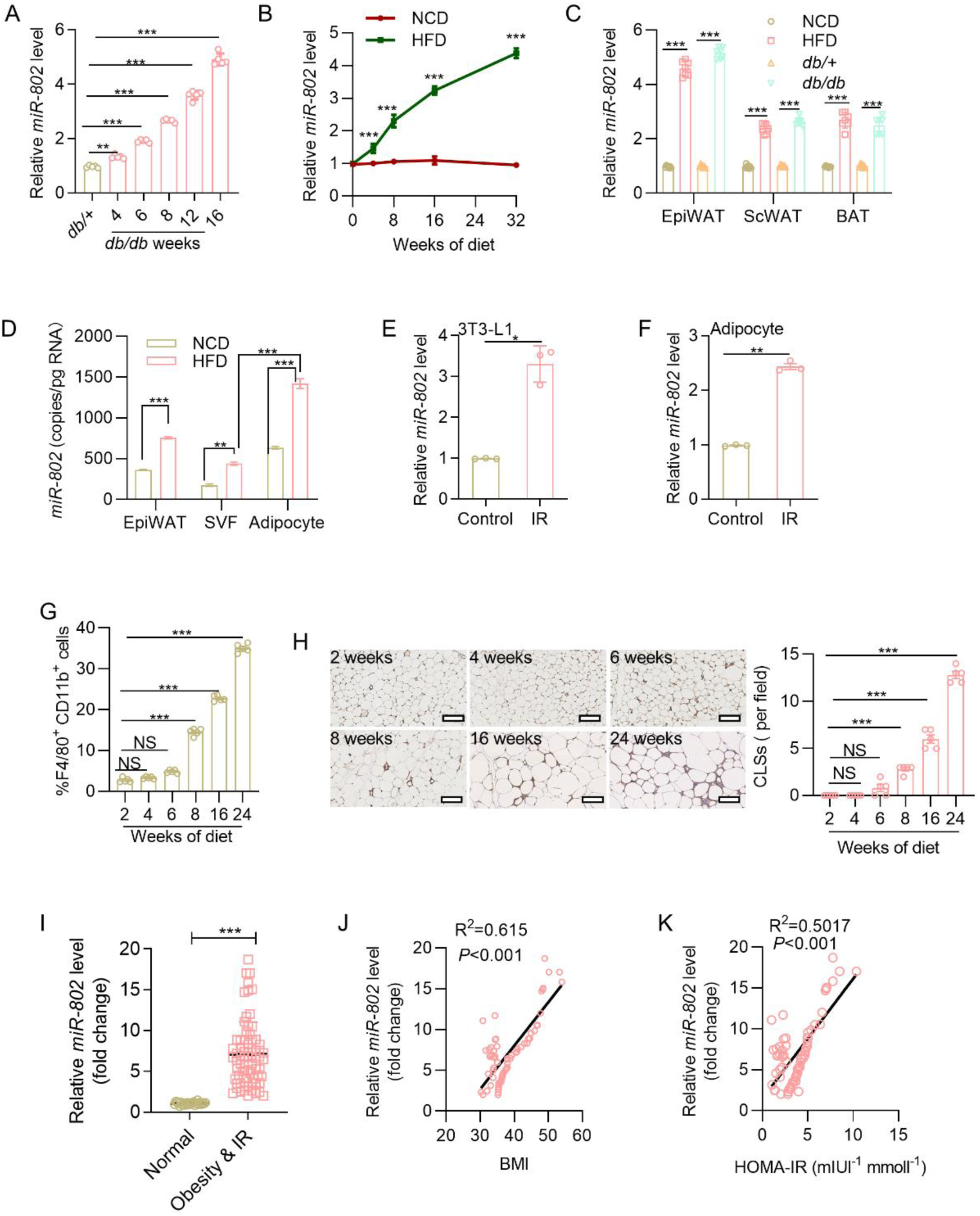
Obesity induced *miR-802* elevation precedes macrophage accumulation. (A) mRNA abundance of *miR-802* in the epiWAT of *db/db* or control mice at 4, 6, 8, 12, and 16 weeks (*n=*5). (B) mRNA abundance of *miR-802* in the epiWAT of mice fed a normal chow diet (NCD) or HFD for 0, 2, 4, 8, 16, 24, and 32 weeks (*n=*5). (C) The expression level of *miR-802* in epiWAT, scWAT and BAT isolated from mice on HFD for 16 weeks or 10 weeks *db/db* mice (*n*=7). (D) Copy number of *miR-802* in mature adipocytes and stromal vascular fraction (SVF) of epiWAT isolated from mice on NCD or HFD for 16 weeks (*n=*5). (E–F) *miR-802* expression levels in insulin resistance 3T3-L1 cell models (E) and insulin resistance WAT SVF cells models (F). (G) F4/80 and CD11b positive cells in SVFs isolated from the epiWAT of mice fed an HFD for 2, 4, 6, 8, 16, and 24 weeks (*n*=5). (H) Representative images of F4/80 staining (left) and quantification of crown-like structures (CLSs; right) in the epiWAT of mice fed an HFD (*n*=5). (I) Expression levels of *miR-802* in human subcutaneous adipose tissue (*n*_normal_=25, *n*_obesity & IR_=70). Scatter plots of *miR-802* expression versus BMI (J) and HOMA-IR (K). Pearson’s correlation coefficients (r) are shown. The fold of *miR-802* was calculated using 2^-ΔΔCt^. Data represent mean ± SEM. *P*-values obtained using a two-tailed unpaired Student’s *t*-test (E, F, I) or two-way ANOVA (A–D, G) are indicated. *\*P*<0.05, *\*\*P*<0.01, *\*\*\*P*<0.001. Relative levels of *miR-802* were normalized to *U6*. epiWAT: epididymal white adipose tissue, scWAT: subcutaneous white adipose tissue, BAT: brown adipose tissue.

Initial studies have indicated that macrophages are responsible for most inflammatory events in adipose tissue (12, 13). However, what initiates macrophage infiltration or the resultant inflammatory cascade is still not well defined. We hypothesized that the elevation of *miR-802* in adipocytes is associated with adipose inflammation and insulin resistance. To test this idea, we examined the correlation between *miR-802* elevation and macrophage infiltration during the progression of diet-induced obesity (DIO). We first carried out a set of flow cytometric analyses to determine the dynamic alterations of macrophages in collagenase-digested SVF from epiWAT. From week 8, the number of double positive CD11b/F4/80 macrophages gradually increased in obese mice (Figure 1G, Figure S1E). Immunohistochemical analysis of F4/80 expression also revealed that the number of macrophages continued to increase in the epididymal fat pads of obese mice as compared to that in mice on a normal diet (Figure 1H). The dynamic increase in *miR-802* preceded the infiltration of macrophages, indicating that *miR-802* may play a critical role in the occurrence of adipose inflammation.

To gain additional insight into the clinical importance of *miR-802* in obese fat, we analyzed the expression of *miR-802* in samples of human subcutaneous adipose tissue. Levels of *miR-802* expression were significantly higher in obese subjects (body mass index [BMI]=38.30±5.82 kg/m^2^, fasting plasma glucose=8.39±1.54 mM, homeostatic model assessment for insulin resistance (HOMA-IR)=3.77±1.97) than in lean ones (BMI=20.55±0.97 kg/m^2^, fasting plasma glucose=4.84±0.53 mM, HOMA-IR=0.21±0.06) (Figure 1I and Figure S1F). Pearson’s correlation analysis showed that the BMI and HOMA-IR were positively associated with *miR-802* abundance in subcutaneous fat (Figure 1J, K). The same phenomenon was also observed in RNA-FISH analysis (Figure S1G), indicating that upregulation of *miR-802* in the adipose tissue during obesity is conserved in humans.

### Adipose-selective overexpression of *miR-802* aggravates inflammatory cascade in obese mice

To further assess the role of adipocyte *miR-802*, we generated adipose-selective *miR-802* KI mice by crossing *miR-802*^ki/ki^ mice (27) with animals expressing Cre recombinase under the control of the promoter of *adiponectin* (Figure S2A, B). Real-time PCR analysis confirmed that the overexpression of *miR-802* was restricted in the adipose tissues of the *miR-802* KI mice, *miR-802* expressions were up-regulated about 150 times, whereas its expression in other organs was not affected (Figure S2C), and the upregulation of *miR-802* was limited to adipocytes and were not observed in SVFs (Figure S2D). There was no obvious difference in food intake, body weight, glucose content, and adiposity between *miR-802* KI mice and their WT littermates in both male and female when they were fed with NCD (Figure S2E-H). We then fed the mice an HFD and performed metabolic and histological analyses. We detected the presence of adipose inflammation, typified by macrophage crown-like structures (CLSs) in epiWAT at 8 weeks in *miR-802* KI mice, which was earlier than their WT littermates, and the number of CLSs was almost doubled at 16 weeks (Figure 2A). No change of CLSs was between two groups fed with NCD (Figure S2I). Consistently, flow cytometric analysis showed that HFD-induced elevation in the number of CD11b^+^F4/80^+^ macrophages in the SVF of epiWAT in adipose-specific *miR-802* KI mice was significantly higher than that in WT littermates in both male and female (Figure 2B and Figure S2J). In *miR-802* KI mice fed on HFD for 16 weeks, the number of classically activated proinflammatory M1 macrophages (defined as CD86^+^CD206^-^) was significantly higher than that of alternatively activated anti-inflammatory M2 macrophages (defined as CD86^-^CD206^+^) in epiWAT (Figure 2C and Figure S2K). In line with this finding, epiWAT from dietary-obese *miR-802* KI mice exhibited obviously higher mRNA expression of the M1 macrophage–related genes (*Ccl2*, *Il-1β, Il-6, Tnf-α, Inos,* and *Ifn-γ*) but significant reductions of M2 macrophage–related genes (*Il-10, Ym1, Arg1*, and *Fizz1*) (Figure 2D). Similarly, HFD also increased the level of several inflammatory factors (chemokine ligand 2 [CCL2], interleukin [IL]-1β, IL-6, and tumor necrosis factor [TNF]-α) in the serum of *miR-802* KI mice (Figure 2E-H).

**Figure 2.**
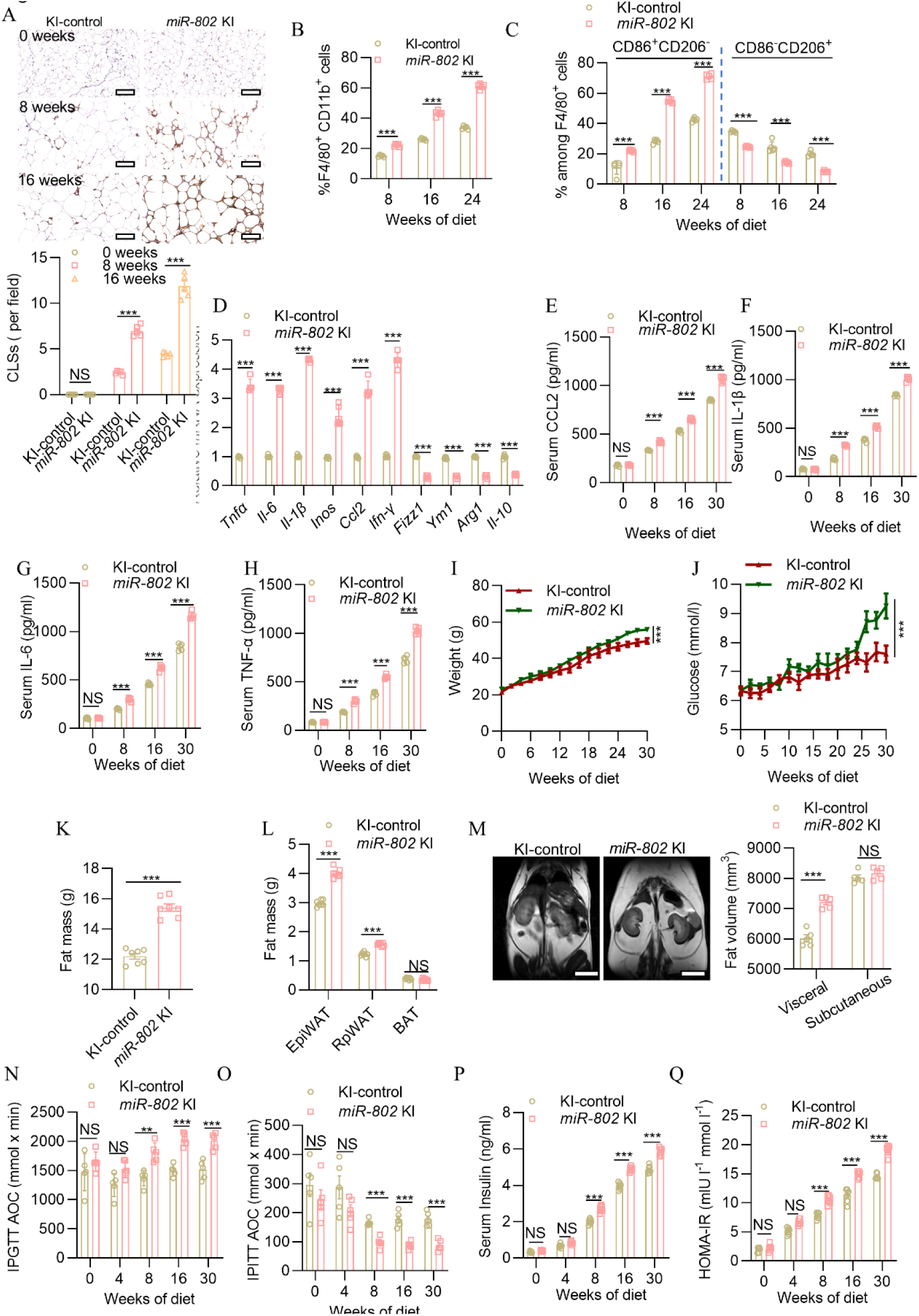
Adipose tissue-specific overexpression of *miR-802* exacerbates adipose tissue inflammation and leads to metabolic dysfunction. (A) Representative images of F4/80 staining (top) and quantification of CLSs (bottom) in epiWAT of WT or *miR-802* KI mice on HFD for 0, 8, and 16 weeks (*n*=5). Scale bar: 40 μm. (B) Percentage of F4/80^+^/CD11b^+^ total macrophages in the epiWAT of *miR-802* KI and KI-control mice fed with HFD (*n*=5). (C) M1 (CD86^+^CD206^-^) and M2 (CD206^+^CD86^-^) within the macrophage population (*n*=5). (D) qRT-PCR analysis for mRNA levels of the M1 and M2 markers in the epiWAT of mice on KI-control or *miR-802* KI at 16 weeks (*n*=5). (E–H) Serum levels of CCL2 (E), IL-1β (F), IL-6 (G), and TNF-α (H) of *miR-802* KI and control mice fed with HFD for 0, 8, 16, and 30 weeks (*n*=5). (I, J) Dynamic changes in body weight (I) and glucose (J) in WT and *miR-802* KI mice during 30 weeks of HFD feeding (*n*=5). (K, L) Fat mass of whole body (K) and individual tissues (L) (*n=*7). (M) Representative coronal section MRI images and visceral and subcutaneous adipose tissue volume of HFD-fed control and *miR-802* KI mice (*n=*5). (N, O) Area over the curve (AOC) of the blood glucose level was calculated via intraperitoneal glucose tolerance tests (IPGTTs, 2 g/kg, N, *n*=5) or intraperitoneal insulin tolerance tests (IPITTs; 0.75 U/kg, O, *n*=5). (P) Fasting insulin (FINS) levels of HFD-fed mice were measured by ELISA (*n*=7). (Q) HOMA-IR was calculated with the equation FBG (mmol l^−1^) × FINS (mIU l^−1^))/22.5. Data represent mean ± SEM. Differences between groups were determined by ANOVA (B–J, L, and N–Q) or two-tailed unpaired Student’s *t*-test (K). *\*P*<0.05, *\*\*\*P*<0.001. Gene levels were normalized to *18S* RNA abundance.

We next explored whether the aggravation of adipose inflammation in adipose-selective *miR-802* KI mice in both male and female were associated with exacerbation of metabolism and insulin sensitivity. We found that in *miR-802* KI mice, HFD induced weight gain (Figure 2I and Figure S2L) and hyperglycemia (Figure 2J) both in male and female. HFD also induced adiposity in *miR-802* KI mice, which mainly manifested in the expansion of visceral WAT (Figure 2K, L). MRI analysis confirmed that HFD induced an increase in visceral WAT in *miR-802* KI mice (Figure 2M). We next monitored the dynamic changes in insulin sensitivity at different time points (0, 4, 8, 16, and 30 weeks) after feeding the two groups of mice with an HFD. As expected, *miR-802* KI mice on a HFD exhibited progressive development of glucose intolerance (Figure 2N and Figure S2M-Q) and insulin resistance (Figure 2O and Figure S2R-V) at 8 weeks, as compared to their WT littermates. These differences became even more obvious after 16 and 30 weeks, coupled with an increase in fasting insulin levels (Figure 2P) and HOMA-IR (Figure 2Q). Collectively, these effects of adipose-selective overexpression of *miR-802* show that *miR-802* is required for the recruitment of macrophages into obese adipose tissue and for the initiation and propagation of the inflammatory cascade.

### *miR-802* depletion ameliorates obesity-induced metabolic dysfunction

Given the striking effects of adipose-selective overexpression of *miR-802* on metabolism, we next investigated whether selectively ablated *miR-802* in adipose tissue could improve metabolic disturbance and inflammation induced by obesity. We generated *miR-802* conditional knockout mice using the Cre/Lox system (Figure S3A). *miR-802*^fl/fl^ were crossed with *Adipoq*-Cre transgenic animals to selectively ablate *miR-802* in adipose tissues (Figure S3B). Expression analysis showed that total *miR-802* levels were reduced by approximately 70% in the adipose tissue but not in SVFs of *miR-802* KO mice compared with WT littermates (Figure S3C, D). The knockout of *miR-802* in adipose tissue did not alter food intake, body weight, glucose level, and adiposity (data not shown); however, this approach could prevent HFD-induced weight gain and hyperglycemia (Figure 3A, B and Figure S3E). Adipose-selective ablation of *miR-802* also alleviated HFD-induced adiposity, mainly by reducing the expansion of visceral WAT, including epiWAT and retro-peritoneal WAT (Figure 3C, D). MRI analysis confirmed this result (Figure 3E). Histological and FACS analysis showed that *miR-802* depletion reduced macrophage infiltration, which mainly manifested as a decrease in the number of CLSs and macrophages (Figure 3F, G and Figure S3F), but had little effect between two groups fed with NCD (Figure S3G). Notably, the *miR-802* KO mice exhibited obvious reductions in mRNA expression of the M1 macrophage-related genes (*Ccl2*, *Il-1β, Il-6, Tnf-α, Inos,* and *Ifn-γ*) but significant upregulation of M2 macrophage-related genes (*Fizz1, Ym1, Arg1*, and *Il-10*) (Figure 3H). The *miR-802* KO mice also markedly blunted HFD-induced elevation in serum levels of several inflammatory factors (TNF-α, IL-6, IL-1β, and CCL2) (Figure 3I). In addition, the insulin resistance and glucose intolerance induced by an HFD were ameliorated by *miR-802* depletion (Figure 3J, K and Figure S3H-M). These phenomena were the same both in male and female *miR-802* KO mice.

**Figure 3.**
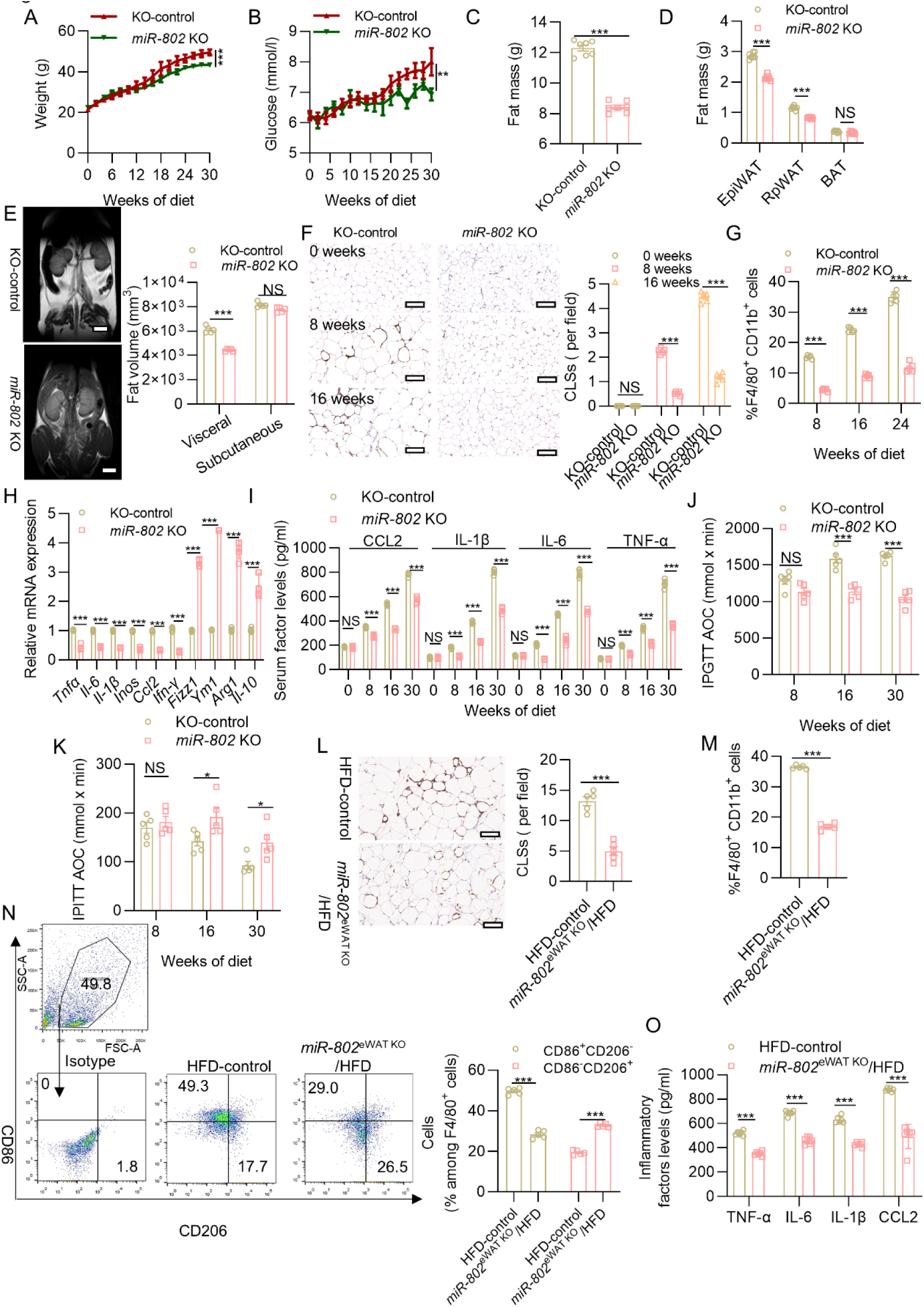
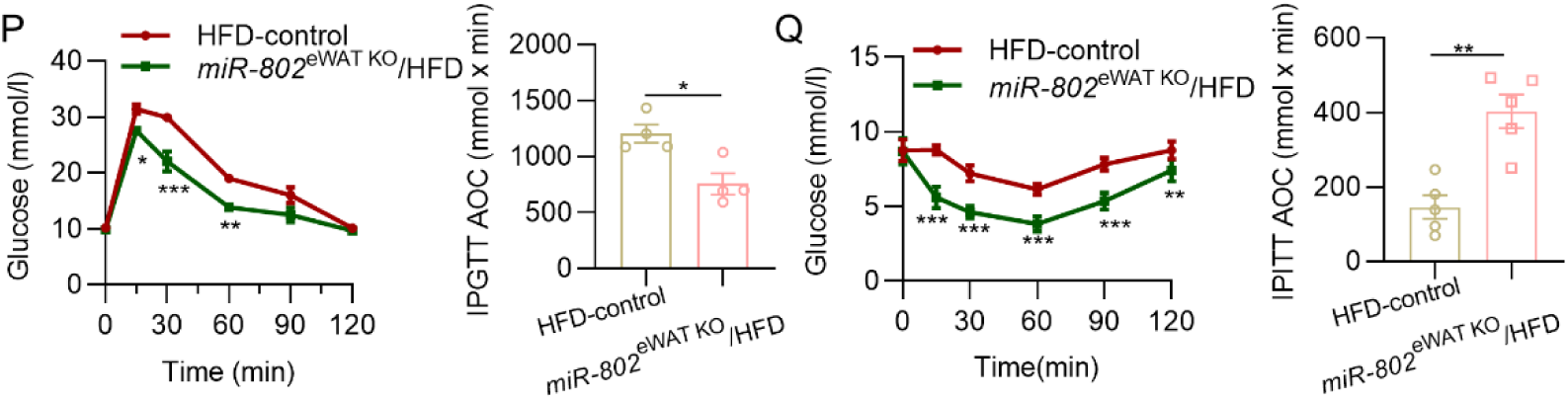
Adipose tissue–specific ablation of *miR-802* protects mice from obesity-induced metabolic dysfunction. (A-B) Dynamic changes in body weight (A) and glucose (B) of KO control and *miR-802* KO mice during 30 weeks of HFD feeding (*n*=7). (C-D) Fat mass of whole body (C) and individual tissues (D) (*n*=7). (E) Representative coronal section MRI images and visceral and subcutaneous adipose tissue volume of HFD-fed control and *miR-802* KO mice (*n*=5). (F) Representative images of F4/80 staining (left) and quantification of CLSs (right) in epiWAT of WT or *miR-802* KO mice on HFD for 0, 8, and 16 weeks (*n*=5). Scale bar: 40 μm. (G) Cells isolated from SVFs of epiWAT in *miR-802* KO and WT mice fed with HFD for 8, 16, and 24 weeks were subjected to flow cytometry analysis for percentage of CD11b^+^/F4/80^+^ total macrophages (*n*=5). (H) qRT-PCR analysis for the mRNA levels of the M1 and M2 markers in epiWAT of mice on HFD 16 weeks (*n*=5). (I) Serum levels of CCL2, IL-1β, IL-6, TNF-α determined with ELISA (*n*=5). (J-K) AOC of the blood glucose level was calculated via IPGTT (1.5 g/kg, J, *n*=5) or IPITT (0.75 U/kg, K, *n*=5). (L) Representative images of F4/80 staining and quantification of CLSs (*n*=5) in the epiWAT of WT or *miR-802* KO mice. Scale bar: 40 μm. (M-N) The percentage of CD11b^+^/F4/80^+^ total macrophages (M, *n*=5) and M1 (CD86^+^CD206^-^), and M2 (CD206^+^CD86^-^) within the macrophage population (N, *n*=5) in the SVFs isolated from epiWAT in the HFD-control or *miR-802*^eWAT^ ^KO^/HFD mice. (O) Serum levels of TNF-α, IL-6, IL-1β, CCL2 determined with ELISA (*n*=6). (P-Q) IPGTT (P) and IPITT (Q) were performed in HFD-control mice or *miR-802*^eWAT^ ^KO^/HFD mice(*n*=5). Data represent mean ± SEM. Differences between groups were determined by ANOVA (A-B, D, E-K, N-Q) or two-tailed unpaired Student’s *t* test (C, L-M). *\*\*\*P* < 0.001. Gene levels were normalized to *18S rRNA* abundance.

We next examined the activity of *miR-802* in obese adipose tissues in which inflammation had already been established. We performed the acute deletion of adipocyte *miR-802* that did not influence whole-body weight. To address this question, we depleted *miR-802* in eWAT using an approach of adeno-associated virus (AAV, *miR-802*^eWAT^ ^KO^) to 16-week-old DIO mice that had been fed an HFD since they were 4 weeks old (Figure S3N). After 7 days, we detected 70% lower expression of *miR-802* compared with the control in the epididymal fat pad; *miR-802* expression was unaffected in other organs (Figure S3O). The weight and the number of CLSs were lowered with *miR-802* sponge treatment (Figure S3P, Figure 3L), and the reduction in macrophage infiltration was confirmed by CD11b and F4/80 flow cytometry analysis (Figure 3M, Figure S3Q). Phenotypic analysis indicated that *miR-802* inhibitor also lowered the M1 (CD86^+^CD206^-^) macrophage fraction, while it increased the M2 macrophage (CD206^+^CD86^-^) fraction (Figure 3N). DIO led to upregulated mRNA expression of proinflammatory cytokines (IL-1β, IL-6, and TNF-α) in the adipose tissue that was suppressed in the *miR-802*^eWAT^ ^KO^ mice (Figure 3O). *miR-802* inhibitor treatment also ameliorated insulin resistance and glucose intolerance in DIO mice (Figure 3P, Q). These results clearly show that *miR-802* inhibitor treatment suppresses preexisting adipose inflammation, which strongly suggests that *miR-802* is required for the maintenance of inflammatory reactions in obese adipose tissue.

### Interplay between *miR-802* ectopically expressed adipocytes and macrophages

We next analyzed the cellular interplay via which inflammation develops in obese adipose tissue. Based on the findings of the *in vivo* experiments summarized above, we hypothesized that obese adipose tissue upregulates *miR-802*, and *miR-802-* overexpressing adipocytes in turn recruit and activate macrophages. To test this hypothesis, we first co-cultured isolated primary macrophages with WAT SVF cells isolated from lean or obese mice to determine whether obese adipose tissue can affect macrophages (Figure 4A). EdU assays and flow cytometric analysis showed that obese WAT SVF induced the proliferation of macrophages, whereas lean fat did so only mildly (Figure S4A, B). Transwell co-culture further showed that obese WAT SVF also promoted macrophage migration and invasion (Figure 4B). We next explored the effects of obese WAT SVF on the characteristics of macrophages. After co-culture macrophages and WAT SVF of obese mice, isolated primary macrophages had elevated expression of classical activation (M1-like) marker CD86, whereas the alternative activation marker (M2-like) CD206 was decreased (Figure 4C). The results of ELISA indicated that obese WAT SVF-induced macrophages were predominantly polarized to pro-inflammatory macrophages (Figure 4D). When we plated primary macrophages in Boyden chambers and treated them with a medium conditioned with obese WAT SVF or lean WAT SVF, the number of macrophages that migrated through the pores between chamber wells with obese WAT SVF conditioned medium was significantly higher than the number of cells cultured in lean WAT SVF conditioned medium (Figure 4E). ELISA results showed that conditioned medium of obese WAT SVF can secrete more humoral factors known to induce macrophage migration, especially CCL2 (Figure 4F).

**Figure 4.**
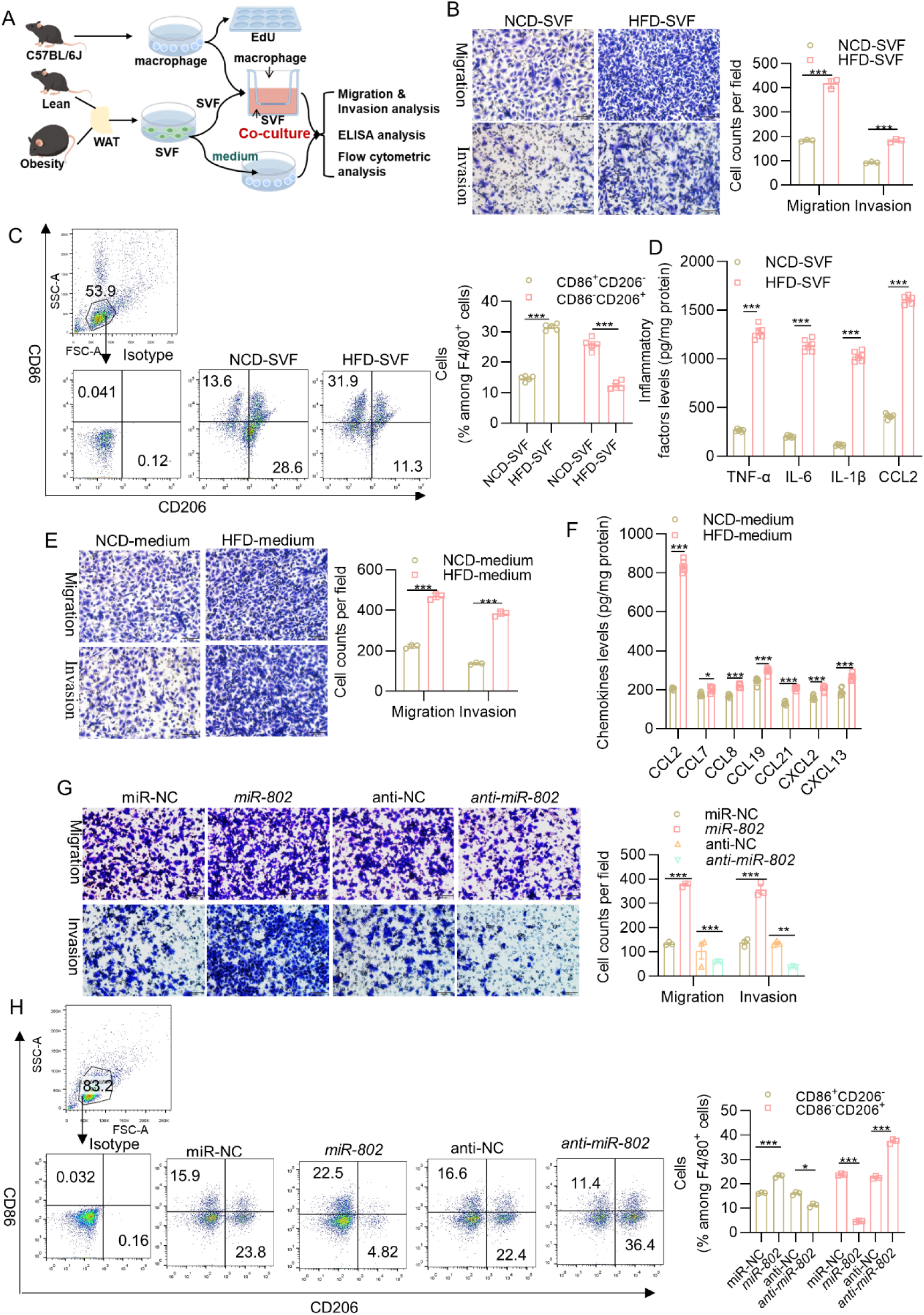
Interplay between miR-802 ectopically expressed adipocytes and macrophages. (A) Flowchart of the co-culture experiments designed for determining WAT SVF of obese adipose tissue can affect macrophages. (B) Obesity promoted macrophage migration and invasion in transwell migration and invasion assay. (C) M1 (CD86^+^CD206^-^) and M2 (CD206^+^CD86^-^) within the macrophage population. (D) The levels of TNF-α, IL-6, IL-1β, and CCL2 determined with ELISA. (E) Migration and invasion ability of macrophages treated with a medium conditioned with obese or lean SVF cells. (F) Chemokine levels in the medium conditioned with obese or lean SVF cells. (G) *miR-802* induced 3T3-L1 cells recruitment more RAW 264.7 cells in transwell migration and invasion assay. (H) *miR-802* mimics-transfected 3T3-L1 cells promoted RAW 264.7 cells M1-like polarization. Data represent mean ± SEM. Differences between groups were determined by ANOVA (D, F). *\*\*P*<0.01, *\*\*\*P*<0.001.

To further confirm the function of *miR-802* in adipose tissue, the adipocyte cell line 3T3-L1 was transfected with *miR-802* mimics (*miR-802*) or *miR-802* inhibitor (*anti-miR-802*). We then explored the effect of *miR-802* ectopically expressed 3T3-L1 cells on the macrophage cell line RAW 264.7 in co-culture. The knockdown and overexpression efficiencies were approximately 80% and 240-fold, respectively (Figure S4C). First, we found that *miR-802*-overexpressing 3T3-L1 cells had no effect on the proliferation and lipid droplet production of RAW 264.7 cells (Figure S4D, E, F). However, *miR-802*-overexpressing 3T3-L1 cells promoted the migration and invasion of RAW 264.7 cells, whereas 3T3-L1 cells knocked down by *anti-miR-802* had the opposite effect (Figure 4G). *miR-802* mimics-transfected 3T3-L1 cells also promoted RAW 264.7 cells M1-like polarization (Figure 4H). We also found higher level of CCL2 in the medium conditioned with *miR-802*-overexpressed 3T3-L1 cells (Figure S4G). Collectively, the results of the co-culture experiments showed that the interaction between *miR-802* ectopically expressed adipocytes and macrophages is crucial for the initiation and propagation of adipose tissue inflammatory cascades.

### *miRNA-802* promotes adipose tissue inflammation and insulin resistance by targeting TRAF3

To better understand the role of *miR-802* in regulating macrophage-mediated adipose tissue inflammation and insulin resistance, we next set out to identify the target genes of *miR-802* in adipocytes. For that, we utilized RNA-sequencing of samples derived from the epiWAT of *miR-802* KI mice and their WT littermates. A total of 191 differentially expressed genes were identified. The cutoff criteria for significant differentially expressed genes were log fold change > 2 and adjusted *p* value < 0.05. We identified 29 upregulated genes and 57 downregulated genes (Figure 5A left, Supplementary Table 2). Then, we combined the multiMiR database(29) with prediction programs (TargetScan Release 7.0 and miRPathDB) to predict possible targets of *miR-802*. Among 18 tested potential targets, TNF receptor-associated factor 3 (*Traf3*) was identified as a genuine target of *miR-802*, which was among the genes that were significantly downregulated in *miR-802* KI versus WT epiWAT (Figure 5A right, Figure S5A). Indeed, we observed that TRAF3 was decreased in both mRNA and protein levels in obese humans and in various obese mice (Figure 5B, C and Figure S5B, C). The targeting potential between *miR-802* and *Traf3* was also observed in *miR-802* KI and *miR-802* KO mice (Figure 5D and Figure S5D). We then demonstrated *miR-802* binding to the *Traf3* 3’-UTR by transiently co-expressing luciferase reporter fusions of *Traf3* and *miR-802* mimics in 3T3-L1 cells. The results of these co-transfection experiments indicated that the relative luciferase activity in *Traf3* 3’-UTR-expressing cells was significantly inhibited by *miR-802*, whereas other *Traf3* 3’-UTR fusions that contained mutations (*Traf3*-MUT) in *miR-802* binding sites were unaffected (Figure 5E). Consistent with these findings, the ectopic expression of *miR-802* in 3T3-L1 cells effectively regulated the mRNA and protein levels of endogenous *Traf3* (Figure 5F). Moreover, we conducted anti-Ago2 RIP in 3T3-L1 cells, which transiently overexpressed *miR-802*. Endogenous *Traf3* pulldown by Ago2 was specifically enriched in *miR-802*-transfected cells (Figure 5G) and vice versa (Figure S5E). Overall, these data suggest that *Traf3* is a direct target of *miR-802*.

**Figure 5.**
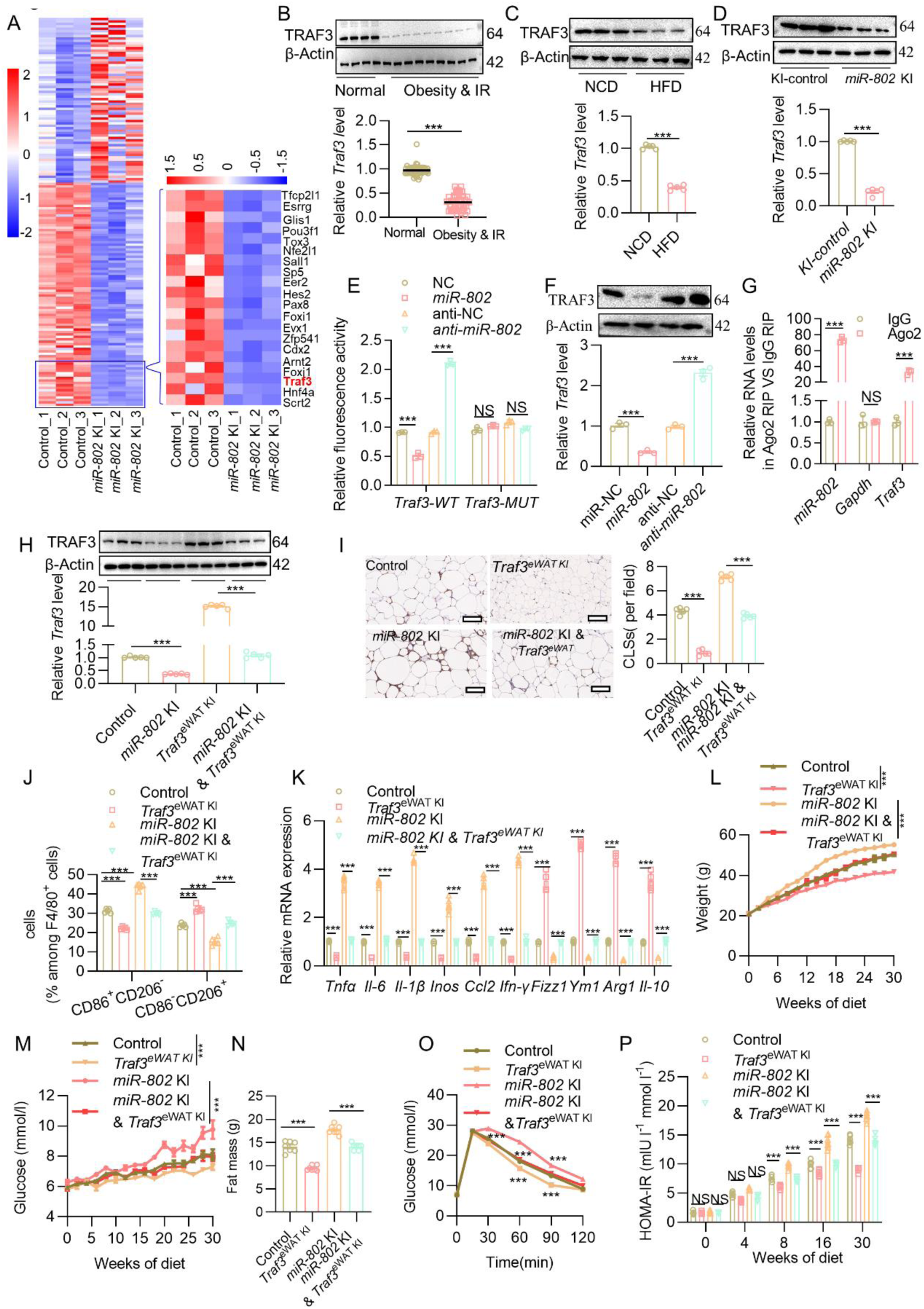
Adipose *miR-802* modulates infiltration and polarization of macrophages by directly targeting *Traf3*. (A) Heat map illustrating the differential expression of mRNAs in the epiWAT of *miR-802* KI mice compared to their WT *miR-802*^ki/ki^ littermates (*n*=3). (B) mRNA and protein levels of TRAF3 in human subcutaneous adipose tissues from obese and normal individuals (*n*_normal_=4 and *n*_obesity&IR_=9). (C, D) mRNA and protein levels of TRAF3 in the epiWAT of HFD mice (C, *n*=3) or *miR-802* KI mice (D, *n*=3). (E) Relative luciferase activity in 3T3-L1 cells co-transfected with *miR-802* mimics and a luciferase reporter containing either *Traf3*-*WT* or *Traf3*-*MUT*. Data are presented as the relative ratio of Renilla luciferase activity to firefly luciferase activity. (F) mRNA and protein levels of TRAF3 in 3T3-L1 cells transfected with *miR-802* mimics or *miR-802* inhibitor. (G) Anti-Ago2 RIP was performed in 3T3-L1 cells transiently overexpressing *miR-802*, followed by qRT-PCR to detect *Traf3* associated with Ago2 (nonspecific IgG served as a negative control). (H) mRNA and protein levels of TRAF3 in the epiWAT of control, *miR-802* KI, *Traf3^eWAT^ ^KI^*, and *miR-802* KI & *Traf3^eWAT^ ^KI^*mice (*n*=3–5). (I) Representative images of F4/80 staining and quantification of CLSs (*n*=5). (J) M1 (CD86^+^CD206^-^) and M2 (CD206^+^CD86^-^) within the macrophage population (*n=*5). (K) qRT-PCR analysis of the mRNA levels of M1 and M2 markers in the epiWAT of HFD-fed control, *Traf3^eWAT^ ^KI^*, *miR-802* KI, and *miR-802* KI & *Traf3^eWAT^ ^KI^*(*n=*5). (L, M) Dynamic changes in body weight (L), glucose level (M), fat mass (N), glucose tolerance (O), and HOMA-IR (P) of control, *miR-802* KI, *Traf3^eWAT^ ^KI^*, and *miR-802* KI & *Traf3^eWAT^ ^KI^*mice during 30 weeks of HFD feeding (*n=*7). Data represent mean ±SEM. Differences between groups were determined by ANOVA (E, F, J–P). *\*\*\*P*<0.001. *MiR-802* abundance was normalized to *U6* level, and other genes levels were normalized to *18S rRNA* abundance.

To address whether the increase in inflammation and insulin resistance in *miR-802* KI mice was attributable to decreased *Traf3*, 8-week-old male *miR-802* KI mice were given AAV-*Adipoq*-*Traf3* (*miR-802 KI & Traf3*^eWAT^ ^KI^) through epididymal fat pad. At 1 week after injection of adeno-associated virus (AAV) expressing *Traf3*, *Traf3* expression in the epiWAT of *miR-802*-KI mice was increased to a level similar to that in WT mice (Figure 5H). Notably, upregulation of *Traf3* led to significant decreases in the counts of total macrophages (Figure 5I and Figure S5F) and M1 macrophages (Figure 5J and Figure S5G) in the epiWAT of HFD-fed *miR-802* KI mice compared with those treated with AAV8-vector. Coherently, the increased expression of M1 macrophage-associated proinflammatory factors (*Tnfα, Il-6, Inos, Il-1β*, and *Ifn-γ*) in the epiWAT of HFD-fed *miR-802* KI mice was reversed by the AAV-mediated upregulation of *Traf3* (Figure 5K). In addition, *Traf3^eWAT^ ^KI^* reversed the weight gain (Figure 5L), hyperglycemia (Figure 5M), and adiposity (Figure 5N) induced by overexpression of *miR-802.* MRI analysis further confirmed that *Traf3* can reverse the increase in visceral fat caused by *miR-802* (Figure S5H). Consistent with these findings, upregulation of *Traf3* led to restoration of glucose intolerance (Figure 5O) and insulin resistance (Figure S5I) after 16 weeks of *Traf3^eWAT^* treatment in HFD-fed *miR-802* KI mice, coupled with a decrease in fasting insulin levels (Figure S5J) and ameliorative HOMA-IR (Figure 5P). Taken together, these findings support the notion that elevated *miR-802* induces macrophage recruitment and polarization at least partly via downregulation of *Traf3*, thereby leading to adipose tissue inflammation and insulin resistance.

### *miR-802* activates noncanonical and canonical NF-κB pathways leading to macrophage recruitment

To further unravel the mechanism by which inhibition of TRAF3 expression induces adipose tissue inflammation, we looked for TRAF3 downstream cascades. Several studies have suggested that TRAF3 negatively regulates the noncanonical NF-κB pathway(30-32), which is consistent with the KEGG analysis based on our RNA-seq results (Figure S6A). This prompted us to measure NF-κB inducing kinase (NIK) protein levels and to explore the processing of p100 to p52. To test whether *miR-802* is required for the suppression of NIK protein levels, western blot analysis of NIK was performed on *miR-802*-overexpressed 3T3-L1 cells and *miR-802* ectopically expressed adipose tissue. As shown in Figure 6A and B, profound accumulation of NIK was observed in all cells with overexpression of *miR-802*, which correlated well with decreased TRAF3. *miR-802* selectively ablated adipose tissues showed the opposite result (Figure S6B). Processing of the p100 precursor to p52, the hallmark of noncanonical NF-κB activation, was also assessed by immunoblotting. Although 3T3-L1 cells exhibited the normal kinetics of p100 processing with substantial p52 accumulation by 48 h after treatment with the empty vector, *miR-802*-overexpressing 3T3-L1 cells and *miR-802* selectively overexpressed adipose tissues showed constitutive and total processing of the p100 precursor protein (Figure 6C, D). On the contrary, there was less accumulation of p52 in *miR-802* selectively deleted adipose tissue (Figure S6C). As expected, IKK-α phosphorylation levels were also enhanced in 3T3-L1 cells and in the epiWAT of *miR-802* KI mice (Figure 6E, F and Figure S6D). To confirm that *miR-802* activates the noncanonical NF-κB pathway through TRAF3, *Traf3* plasmid was transfected into *miR-802*-overexpressing 3T3-L1 cells, then NIK protein levels and processing of p100 to p52 were again assessed by immunoblotting. As shown in Figure 6G, *Traf3* restored the levels of NIK and the processing of p100 to p52 in *miR-802*-overexpressing 3T3-L1 cells. Moreover, these results were confirmed in *miR-802* KI mice (Figure 6H), indicating that *miR-802* regulated the noncanonical NF-κB pathway via TRAF3.

**Figure 6.**
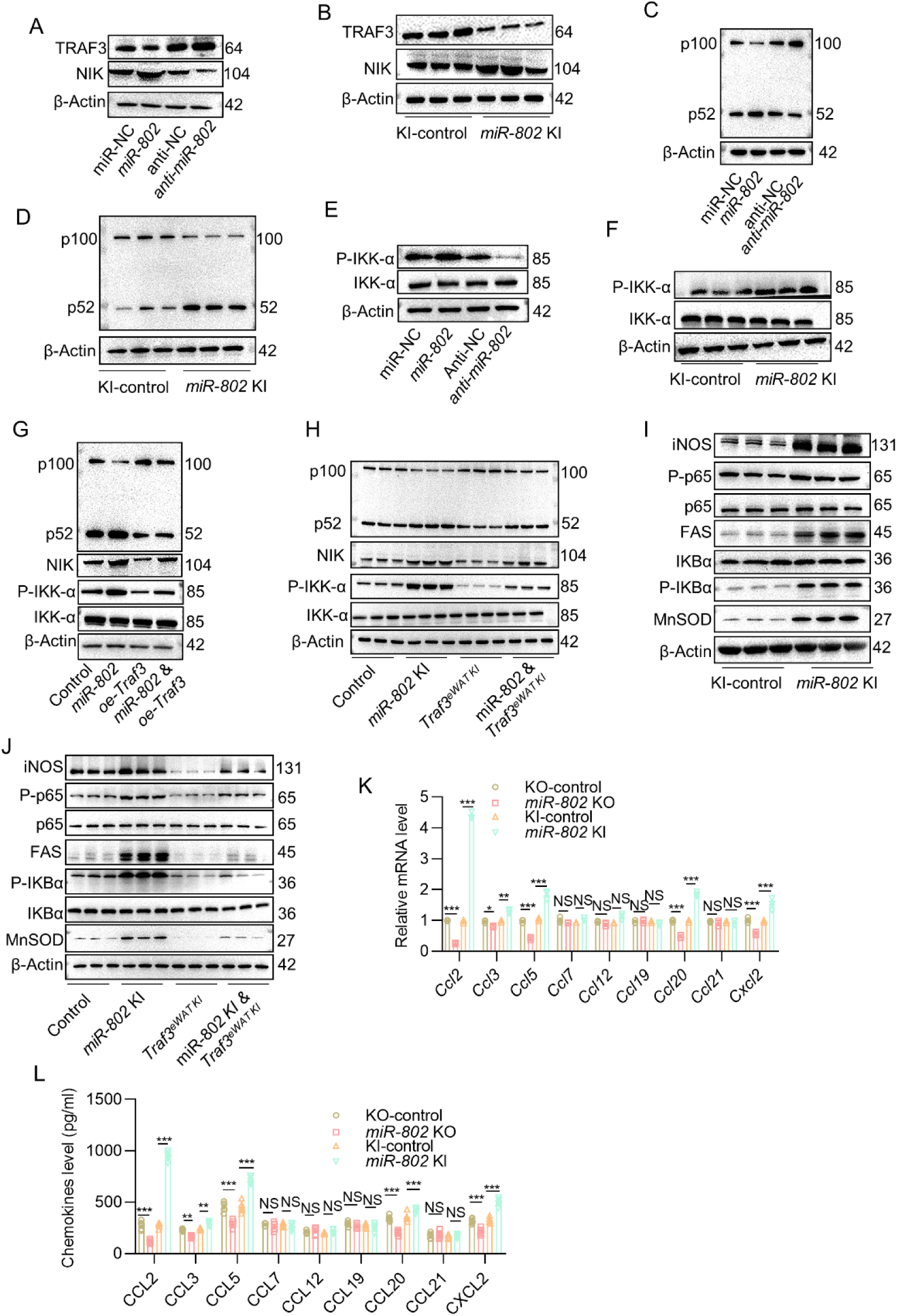
*miR-802* activates noncanonical and canonical NF-κB pathways by recruiting macrophages. (A, B) NIK protein levels in 3T3-L1 cells transfected with *miR-802* mimics or *miR-802* inhibitor (A) in the epiWAT of *miR-802* KI mice (B, *n*=3). (C, D) P100/52 protein levels in 3T3-L1 cells transfected with *miR-802* mimics or *miR-802* inhibitor (C), and in the epiWAT of *miR-802* KI mice (D, *n*=3). (E, F) Protein levels of IKK-α and P-IKK-α in 3T3-L1 cells transfected with *miR-802* mimics or *miR-802* inhibitor (E) in the epiWAT of *miR-802* KI mice (F, *n*=3). (G, H) Overexpression of *Traf3* reverses the protein levels of NIK, P-IKK-α, and P100/52 in 3T3-L1 cells (G) and in the epiWAT of *miR-802* KI mice (H, *n*=3). (I, J) Protein levels of some major canonical NF-κB signaling targets in the epiWAT of *miR-802* KI mice (I, *n*=3) and *Traf3^eWAT^ ^KI^* rescued mice (J, *n*=3). (K, L) qRT-PCR (K) and ELISA (L) were performed to detect major chemokine levels. Data represent mean ± SEM. Differences between groups were determined by ANOVA (K–L). *\*\*\*P*<0.001. Genes levels were normalized to *18S rRNA* abundance.

Previous studies have suggested that TRAF3 also suppresses activation of the canonical NF-κB pathway (33, 34). To verify whether *miR-802* can regulate the canonical NF-κB pathway through TRAF3, nuclear extract was harvested from the adipose tissue of 16-week-old WT and mice with adipose-selective forced expression of *miR-802*. NF-κB activation status was then assessed by measuring p65, IκBα, and some major targets associated with the pathway. As shown in Figure 6I, p65 and IκBα were phosphorylated, and some major targets of canonical NF-κB signaling, such as MnSOD, FAS, and iNOS, were activated in *miR-802* KI mice. As expected, the activation of canonical NF-κB signaling in *miR-802* KI mice was partially reversed by overexpression of *Traf3* (Figure 6J). To assess the potential impact of heightened NF-κB activity, we harvested mRNA from the adipose tissue of WT and *miR-802* KI mice and analyzed the expression levels of multiple noncanonical and canonical NF-κB pathway target genes(35, 36) using qRT-PCR. Here, we observed that the expression levels of *Ccl2*, *Ccl3*, *Ccl5*, *Ccl20*, and *Cxcl2* were elevated in the adipose tissue of *miR-802* KI mice (Figure 6K), which is consistent with ELISA results using the serum of *miR-802* KI mice (Figure 6L). Taken together, these data indicate that *miR-802* activates the noncanonical and canonical NF-κB pathways via TRAF3, leading to macrophage recruitment.

### *miR-802* promotes lipid synthesis and M1 macrophage polarization in adipose tissue through activating SREBP1

The mechanism of an *miR-802* increasing M1 polarization by inducing NF-κB pathways remains unclear. To better understand the role of *miR-802* in regulating macrophage polarization, we performed transcriptome sequencing using epiWAT derived from *miR-802* KI mice. In the adipose tissues of *miR-802* KI mice, the expression of 191 mRNAs was significantly altered compared to that of mRNAs in WT mice, of which the expression of 75 mRNAs increased (Figure 7A, Supplementary Table 2). We found that seven mRNAs were upregulated more than 10-fold (Figure 7A, right), and these mRNAs were annotated using UCSC and Ensemble. Among these upregulated genes, we focused on the lipogenic gene sterol regulatory element-binding protein 1 a (SREBP1a), which is involved in fatty acid synthesis and lipid droplet formation(37). We verified that the mRNA levels of *Srebp1a* were increased in both the epiWAT of *miR-802* KI mice and in 3T3-L1 cells transfected with *miR-802* mimics in qRT-PCR analysis (Figure S7A and B). We also found that the mature form of the SREBP-1 protein (m-SREBP1) was significantly higher in the epiWAT of *miR-802* KI and 3T3-L1 cells transfected with *miR-802* mimics (Figure 7B and C). As expected, overexpression of *Traf3* reduced the upregulation level of mature SREBP1 induced by *miR-802* (Figure 7D). However, target gene prediction algorithms as well as luciferase reporter and Ago2-RIP assays confirmed that *Srebp1a* is not the direct target gene for *miR-802* (data not shown). This prompted us to verify whether *Srebp1a* is a downstream gene in the NF-κB pathway.

**Figure 7.**
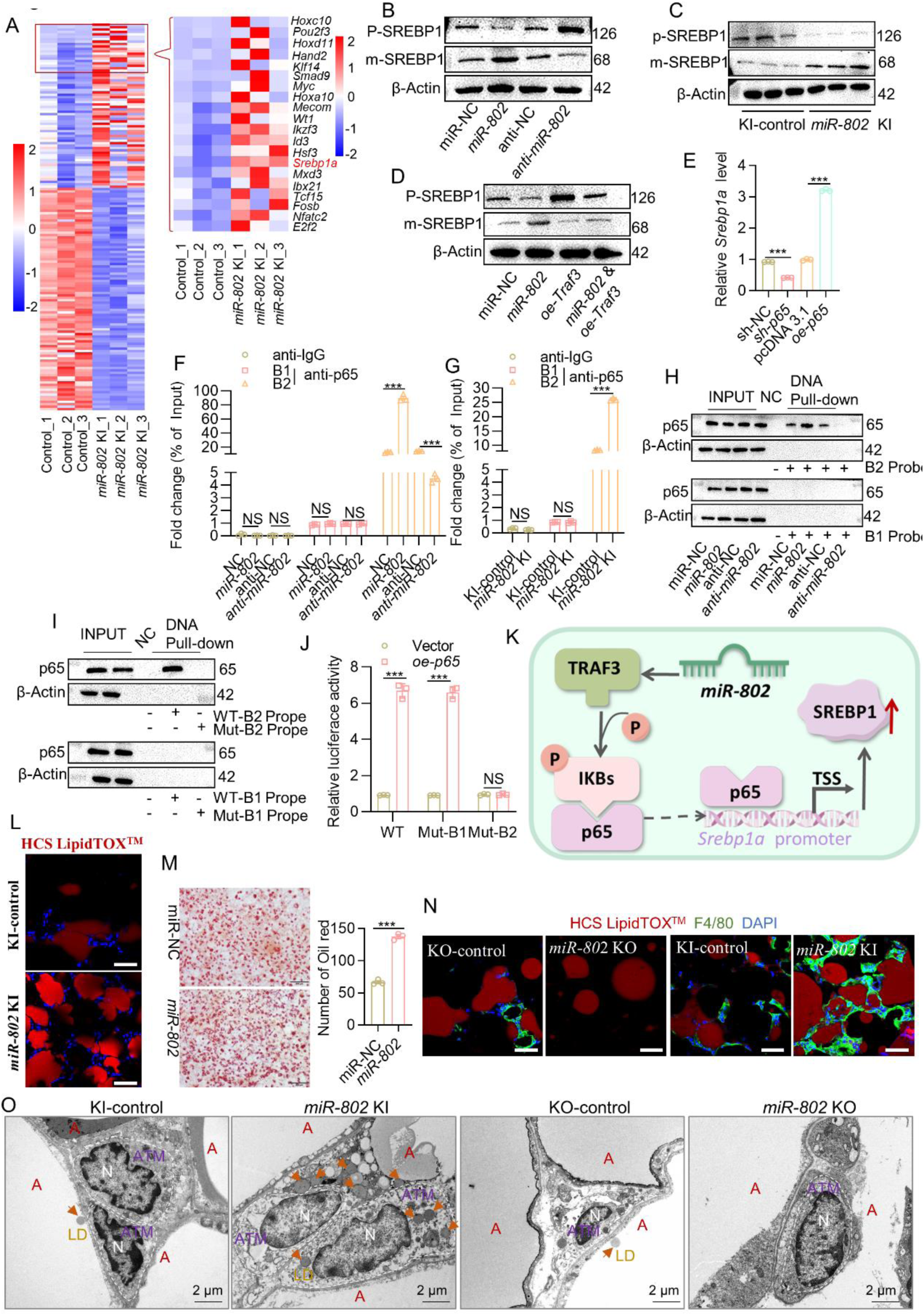
*miR-802* promotes lipogenesis and induces M1 macrophage polarization in adipose tissue through activating SREBP1. (A) Heat map illustrating the top 20 upregulated mRNAs in the epiWAT of *miR-802* KI mice compared to their WT *miR-802^fl/fl^* littermates (*n*=3). (B–C) Protein levels of the mature form of SREBP-1 protein (m-SREBP1) and the precursor form of SREBP-1 (P-SREBP1) in mature 3T3-L1 cells transfected with *miR-802* mimics or *miR-802* inhibitor (B), in the epiWAT of *miR-802* KI mice (C, *n*=3). (D) The protein levels of m-SREBP1 and P-SREBP1 were reversed by *Traf3*. (E) *Srebp1a* mRNA levels in 3T3-L1 cells transfected with *p65*-overexpressing plasmid or *p65* shRNA plasmid. (F, G) ChIP-qPCR assays were conducted to verify that *p65* binds to the *Srebp1* promoter in 3T3-L1 cells transfected with *miR-802* mimics or *miR-802* inhibitor (F) and in the epiWAT of *miR-802* KI mice (G, *n*=3). (H) DNA pull-down assay using a biotinylated DNA probe corresponding to the −360 to −400 or −1198 to −1237 region of the *Srebp1* promoter in 3T3-L1 cells transfected with *miR-802* mimics or *miR-802* inhibitor. (I) DNA pull-down assay using a biotinylated DNA probe corresponding to the −1198 to −1237 region of the wild-type (WT) or a mutant sequence of the *Srebp1* promoter in 3T3-L1 cells stimulated with *p65* plasmid for 48 h. (J) Luciferase reporter assays in 3T3-L1 cells transfected with the indicated plasmids for 48 h. Dual-luciferase activity was determined. (K) Schematic illustration for the mechanism of *miR-802* increased *Srebp1* expression by activating canonical NF-κB pathways. (L) Representative images of the immunofluorescence of lipid droplets (HCS LipidTOX^TM^, Red) and DAPI (Blue). Scale bar: 20 μm. (M) Oil red O staining was performed to assess the number of lipid droplets in 3T3-L1 cells transfected with *miR-802* mimics. Scale bar: 200 μm. (N) Representative images of the immunofluorescence of lipid droplets (HCS LipidTOX^TM^, Red) and F4/80 (Green, *n*=3). Scale bar: 20 μm. (O) Transmission electron microscopy (TEM) was performed to detect the contact between lipid droplets and macrophages (*n*=3). Data represent mean ±SEM. Differences between groups were determined by ANOVA (E–G and J). *\*\*\*P*<0.001. Genes levels were normalized to *18S rRNA* abundance.

For this purpose, we first predicted the nuclear factor-κB (NF-κB) family (*p65*, *RelB*, *C-rel*, *p50*, and *p52*) sites in the promoter of *Srebp1a* using JASPAR and the Promo database. Two *p65* potential binding sites (B1 and B2) were found in the promoter of *Srebp1a* (Figure S7C). We next overexpressed *p65* in 3T3-L1 cells; qRT-PCR results showed that *p65* could increase *Srebp1a* expression in 3T3-L1 cells (Figure 7E). To determine whether *Srebp1a* is a direct *p65* target gene, ChIP-qPCR assays were used. The results showed that occupancy of *p65* binding site 2 (B2) on the *Srebp1a* promoter was significantly increased in *miR-802*-overexpressing 3T3-L1 cells and *miR-802* selectively overexpressed adipose tissues (Figure 7F and G, Figure S7D and E). We then conducted DNA pull-down assays to examine the binding of *p65* to the *Srebp1a* promoter *in vitro*. We constructed two DNA probes containing −360 to −400 or −1198 to −1237 that contained the predicted binding site 1 (B1) and predicted binding site 2 (B2), respectively, to detect binding to *p65* in nuclear extracts. Similar findings were obtained in that the B2 DNA probe, but not the B1 DNA probe, bound to *p65* in the 3T3-L1 cell line overexpressing *miR-802* (Figure 7H). However, with mutant B2 (agggaatgct, Mut2), DNA pull-down results showed that *p65* could not bind to Mut2 (Figure 7I). Moreover, we constructed a luciferase reporter plasmid containing the *Srebp1a* promoter region from −1295 to +1 WT and two mutant reporter plasmids mutated in −375 to −385 (Mut1) or in −1211 to −1221 (Mut2). Overexpression of *p65* significantly enhanced WT and Mut1, but not Mut2-driven luciferase in 3T3-L1 cells (Figure 7J). Taken together, these results indicated that *miR-802* indirectly stimulates *Srebp1a* expression via the canonical NF-κB signaling pathway (Figure 7K).

As previously described, *Srebp1a* is a well-established regulator of lipid synthesis(37). Accordingly, *miR-802* overexpression significantly increased the number and fluorescence intensity of lipid droplets, which correlated well with increased SREBP1 (Figure 7L-M). Conversely, knockdown of *miR-802* strongly reduced lipid droplet formation in 3T3-L1 cells and *miR-802*-KO mouse adipose tissue (Figure S7F and G). Lipid droplets have been shown to play a crucial role in M1 macrophage polarization (38, 39). Consistent with this, we observed that adipose tissue macrophages (ATMs) of the *miR-802* KI mice could engulf more lipid droplets (Figure 7N, O). The elevated expression of the classical activation marker further indicated that lipid droplets induced the ATMs in *miR-802* KI mice to the pro-inflammatory phenotype (Figure S7H). Altogether, these data show that *miR-802* indirectly regulates lipid droplet formation through SREBP1 and ultimately promotes macrophage M1 polarization.

## Discussion

Macrophage infiltration of adipose tissue has been described in both mice and humans during obesity. However, how lipid-loaded hypertrophic adipocytes send signals to trigger infiltration and alter the polarization of macrophages in obesity remains poorly understood. In this study, we found that *miR-802* endows adipose tissue with the ability to interact with macrophages and regulate the inflammatory cascade. Mechanistically, *miR-802* recruits macrophages and drives the polarization program toward proinflammatory M1 phenotype by targeting the cytoplasmic adaptor protein TRAF3 (Figure 8). Our findings indicate that *miR-802* has essential roles in the initiation and maintenance of adipose tissue inflammation and systemic insulin resistance.

**Figure 8.**
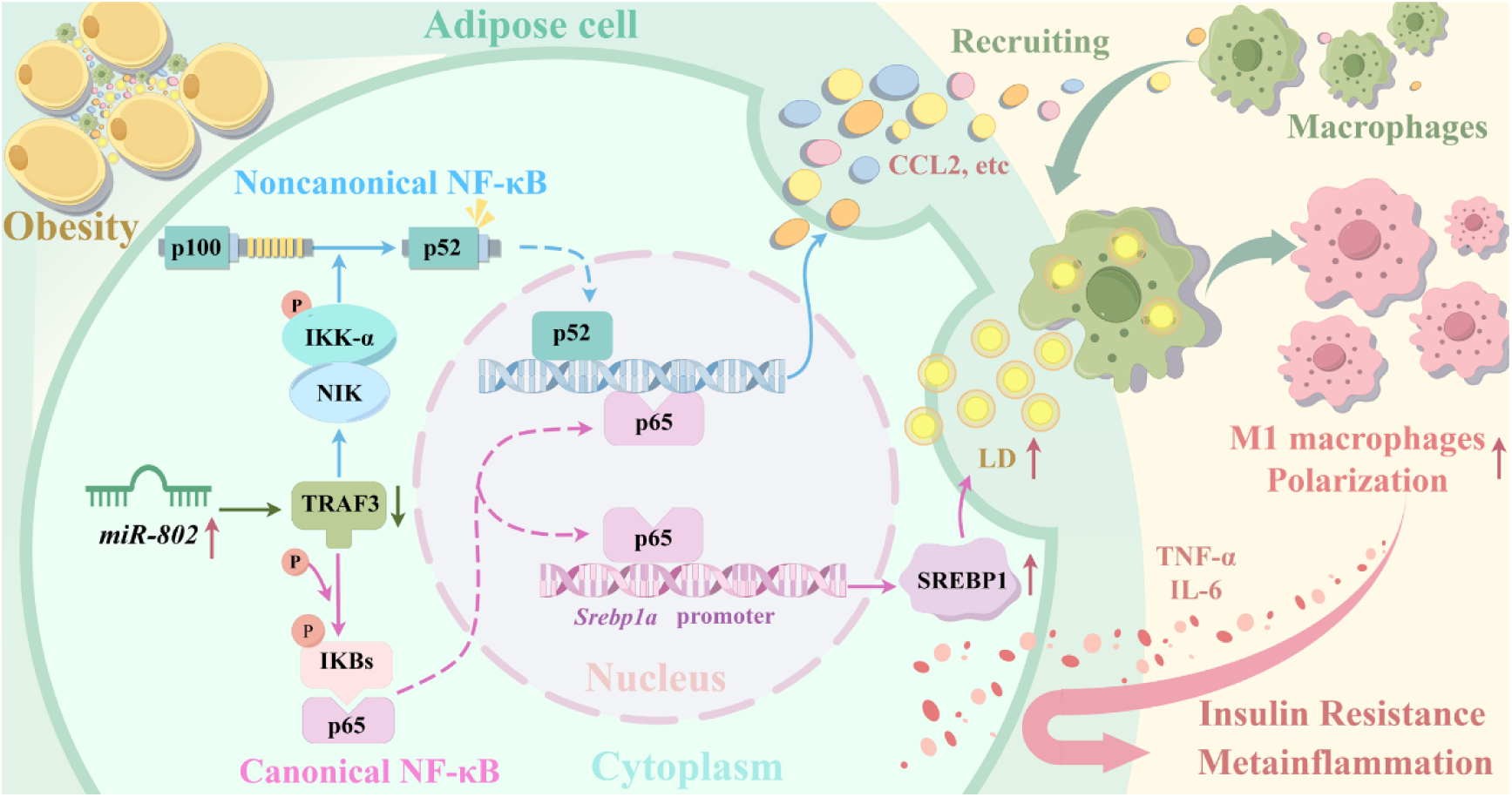
Schematic illustration for the mechanism of *miR-802* exacerbates adipose tissue inflammation and leads to metabolic dysfunction during obesity. We found that *miR-802* endows adipose tissue with the ability to interact with macrophages and regulate the inflammatory cascade. During obesity, *miR-802* promotes adipose tissue secretion more chemokines recruiting macrophages by targeting *Traf3 activating* canonical and noncanonical NF-κB signaling pathways; and *miR-802* increases lipogenesis through promoting *Srebp1* transcription, then, macrophages toward proinflammatory M1 phenotype by engulfing lipid droplet.

Adipose tissue inflammation is a hallmark of obesity and a causal factor of metabolic disorders such as insulin resistance. Mice fed an HFD frequently develop chronic low-grade inflammation within adipose tissues, characterized by increased infiltration of macrophages and the production of pro-inflammatory cytokines. Here, we showed that the increasing trend of *miR-8*02 in adipocytes is an early event during the development of adipose tissue obesity induced by an HFD. *miR-802* expression in visceral fat was progressively increased with the development of dietary obesity, whereas adipose-selective ablation of *miR-802* protected mice from exacerbation of meta-inflammation and insulin resistance caused by dietary stress. The high level of *miR-802* expression in visceral fat may partly explain why this adipose depot is more prone to inflammation and is closely related to insulin resistance. *miR-802* is required for adipose tissue inflammation and has major roles in macrophage recruitment and polarization. Thus, *miR-802* is crucially involved in initiating inflammatory cascades in obese adipose tissue. Moreover, the finding that *miR-802* inhibitor treatment ameliorated pre-established adipose inflammation in DIO mice indicates that *miR-802* is also essential for maintenance of the inflammatory response. Although previously studies have found that *miR-802* was up-regulated in the adipose tissue during obesity (27, 28, 40, 41), while the function of *miR-802* was focused on cancers(42), liver(43, 44), small intestine(41) and pancreas(40). Whether *miR-802* can regulate adipose function is still confused. To our knowledge, the present study is the first to directly address the functional role of *miR-802* in adipose tissue inflammation. The findings that systemic insulin resistance is ameliorated by *miR-802* depletion and is aggravated by adoptive transfer of *miR-802* mimics strongly suggest that *miR-802*-dependent adipose inflammation has an impact on systemic metabolism.

Like most other miRNAs, *miR-802* regulates the expression of multiple genes in different tissues. In the liver, *miR-802* is induced by obesity and impaired glucose tolerance, and it attenuates insulin sensitivity by downregulation of *Hnf1b*(28). Genetic ablation of *miR-802* in the small intestine of mice leads to decreased glucose uptake, impaired enterocyte differentiation, increased Paneth cell function, and intestinal epithelial proliferation through derepression of *Tmed9*(41). We recently discovered that in pancreatic islet cells, elevated *miR-802* causes impaired insulin transcription and secretion by targeting *NeuroD1* and *Fzd5*(27). In this study, we found that *miR-802* promotes adipose tissue inflammation and insulin resistance by targeting TRAF3 in adipocytes. As a member of the TNF receptor (TNFR) superfamily, TRAF3 plays vital roles in inflammatory responses via activation of both the canonical and noncanonical NF-κB signaling pathways(32, 33) following engagement of a variety of TNFR superfamily members such as Baff receptor, lymphotoxin β receptor, and CD40(45). Here, we found that *miR-802* can regulate the NF-κB pathway by directly targeting TRAF3 rather than by activating the classic receptor, which enriches the understanding of the NF-κB pathway.

Macrophage accumulation was significantly higher in adipose tissue from HFD-fed *miR-802* KI mice than in WT mice, suggesting that overexpression of *miR-802* enhances the infiltration ability of macrophages. Correspondingly, we observed that *miR-802-*overexpressing adipocytes released more chemokines by activating NF-κB pathway, such as CCL2, CCL5, CCL20, and CXCL2. Adipose tissue inflammation is well documented as an important contributor to systemic insulin resistance(46). This was further validated by our enhanced adipose tissue inflammatory responses in *miR-802* KI mice. Moreover, HFD-fed *miR-802* KI mice exhibited adipose tissue macrophage infiltration, proinflammatory cytokine expression, and NF-κB pathway activation. Genes that are crucial for meta-inflammation and insulin resistance were directly affected by the enhancement of *miR-802* in adipose tissue. Thus, increased adipose tissue inflammation resulting from *miR-802* overexpression contributed, in large part, to systemic insulin resistance in *miR-802* KI mice.

The chronic inflammation microenvironment is one of the main features of obesity. A recent study found that adipocytes can release lipid-filled vesicles that become a source of lipids for local macrophages(47). Phagocytosis or excessive accumulation of lipid droplets can induce macrophage M1 polarization(47, 48). In our study, we observed the same phenomenon, that is, in *miR-802* KI mice, macrophages accumulated more lipid droplets and exhibited an inflammatory phenotype. SREBP1 has been found to promote the acute inflammatory response and lipogenesis (49, 50). Here, we found that *miR-802* increased SREBP1 expression inducing lipogenesis by activating canonical NF-κB signaling pathways, then macrophage engulf lipid droplet promoting macrophage M1 polarization. This has enriched our understanding of the functionality of SREBP1 to some extent. However, *miR-802* only indirectly regulates SREBP1, but it still has a considerable impact on macrophages, indicating the importance of miRNA positive or indirect regulation.

Taken together, our results support the idea that obese adipose tissue activates *miR-802*, which, in turn, initiates and propagates inflammatory cascades, including the recruitment of macrophages into obese adipose tissues and their subsequent induction of the inflammatory phenotype. Thus, *miR-802* appears to have a primary role in obese adipose tissue inflammation. However, future studies are needed to clarify which environmental cues within obese adipose tissue initiate *miR-802* elevation. The present observations indicate that *miR-802* inhibitors might offer a novel approach to prevent diseases associated with insulin resistance.

## Materials and methods

### Animal studies

All mice used were of mixed strain backgrounds with approximately equal contributions from C57BL/6J, with the exceptions of *db/db* mice (C57BLKS/J). *MiR-802*^fl/fl^ and *miR-802*^ki/ki^ in mice were initially described in(27). To generate adipose-specific *miR-802* knockout and *miR-802* knockin animals, we used *Adipoq*-Cre mice on a C57BL/6J background purchased from Jackson Laboratories. Mice were crossed with homozygous for *miR-802*^fl/fl^ or *miR-802*^ki/ki^ and heterozygous for *Adipoq*-Cre to generate *Adipoq-miR-802* KO mice (*miR-802* KO), *Adipoq*-*miR-802* KI mice (*miR-802* KI), control *miR-802*^fl/fl^ littermates or control m*iR-802*^ki/ki^ littermates. Studies were performed on 8-week-old male and female mice initially housed under standard conditions with full access to standard mouse chow and water. After this time, mice were switched to a 60% high-fat diet (HFD) or normal chow diet (NCD) consisting of a 10% fat diet for 30 weeks. All mice had free access to food and water ad libitum. Animals were housed in a temperature-controlled environment with a12 h dark–light cycle. At the end of the 30-week period, mice were euthanized via overdose of isoflurane anesthesia, and tissues were immediately weighed, dissected, and frozen in liquid nitrogen. Tissue samples were stored at -80 °C until use. Care of all animals was within institutional animal-care committee guidelines, and all procedures were approved by the animal ethics committee of China Pharmaceutical University (Permit Number: 2162326) and were in accordance with the international laws and policies (EEC Council Directive 86/609,1987).

For administration of AAV8-Adipoq*-miR-802* sponge vector, AAV8-Adipoq-*Traf3* vector to epididymal adipose tissue, mice were anesthetized with pentobarbital sodium (60 mg/kg) intraperitoneally and the laparotomy was performed. Each epididymal fat pad was given 8 injections of 5 uL (1 × 10^13^ viral genome copies) of AAV solution.

### Human adipose samples of lean and overweight individuals

Adipose and clinicopathological data were collected from Sir Run Run Hospital, Nanjing Medical University (Nanjing, China). All patients enrolled in this study were obese (BMI > 25). The negative controls were normal-weight individuals (20 ≤ BMI ≤ 25). All human subjects provided informed consent. All human studies were conducted according to the principles of the Declaration of Helsinki and were approved by the Ethics Committees of the Department Sir Run Run Hospital (Nanjing, China, 2023-SR-046). The clinical features of patients are listed in Supplementary Table 1.

### Adipose sample preparation

SVF and mature adipocytes were obtained as follows: adipose tissue samples were digested with collagenase type 1 in Krebs-RingerHenseleit (KRH) buffer for 30 min at 37°C. Cell suspensions containing mature adipocytes and SVF were then filtered with nylon mesh and washed three times with KRH buffer. Mature adipose was floated to the surface and the remaining solution containing the SVF was centrifuged at 1500 rpm for 5 min. The pellet was washed with pre-adipocyte growth medium (DMEM-F12 supplemented with 10% calf serum and 1% penicillin-streptomycin), followed by a second centrifugation. SVF cells were then cryopreserved using a freezing medium (DMEM-F12 supplemented with 60% FBS and 10% DMSO). The medium was added to the pellet and frozen with a temperature gradient (−1°C/minute) and stored in liquid nitrogen until analysis. Following collection, whole adipose tissue samples were quickly frozen in liquid nitrogen and stored until analysis.

### 3T3-L1 cell culture and differentiation

3T3-L1 cells were cultured in DMEM (Gibco) containing 10% calf serum with high glucose at 37 °C, 5% CO_2_ and full saturation humidity until they reached 80%-90% confluence, at which point the media was changed to the first differentiation medium containing high glucose DMEM, 10% FBS, 0.5 mM 3-isobutyl-1-methylxanthine (IBMX), 1 μM dexamethasone and 10 μg/ml insulin for 48 h, then the media was changed to the terminal differentiation cocktail containing high glucose DMEM, 10% FBS, and 10 μg/ml insulin for 48 h.

The insulin-resistant cell models were established in mature 3T3-L1 and mature WAT SVF cells by 0.5 mM palmitate, 10 μg/ml insulin and 25 mM glucose for 24 h.

### Resident peritoneal macrophage isolation and culture

For 2 d, 1.0 ml of sterile 4% thioglycolate medium (Sangon Biotech, China) was injected into the peritoneal cavity of C57BL/6J mice. Then, resident peritoneal macrophages were obtained via peritoneal lavage with 5 ml lavage solution (PBS (Sangon Biotech) supplemented with 5 mM EDTA and 4% FBS). Lavages of the same genotype were pooled and resuspended in complete medium (RPMI 1640 supplemented with 10% FBS, 100 U/ml penicillin, 10 μg/ml streptomycin, and 400 μM L-glutamine [Invitrogen]). Typically, the cells were plated and left to adhere for 3 h at 37°C, 5% CO_2_ before being washed two times with warm complete medium. The cells were plated on transwell permeable supports or 24-well plates and co-cultured with SVF cells.

### Migration and invasion assays

The 3T3-L1 cells or SVF cells were evenly plated in 24-well plates. To differentiate mature cells, migration and invasion assays were performed using a transwell chamber (Millipore, Billerica, MA, USA). For the migration assay, RAW 264.7 macrophage cells were seeded in the upper chamber with serum-free medium (1.0×10^5^ cells); the bottom chamber contained mature 3T3-L1 cells. For the invasion assay, the chamber was coated with Matrigel (BD Biosciences, Franklin Lakes, NJ, USA); the subsequent steps were similar to the migration assay. After the cells migrated or invaded for 24 h, they were fixed and stained with crystal violet. Migrated and invaded RAW 264.7 cells were counted under an inverted light microscope. The number of migrated or invaded cells was quantified by counting the number of cells from 10 random fields at ×100 magnification.

### RNA-sequencing analysis

Total RNA from epididymis white adipose tissue of wide type control mice (*n*=3) and *miR-802* KI mice (*n*=3) was isolated using the RNeasy mini kit (Qiagen) following the protocol. The quality of the samples, the experiment, and the analysis data was completely finished by the HaploX (Shangrao, China). Cuffdiff (v2.2.1) 51 was used to calculate the fragments per kilobase million (FPKM) for mRNAs in each group. A difference in gene expression with a *p* value ≤ 0.05 was considered significant. The raw data is presented in Supplementary Table 2. The RNA-seq raw data that support the findings of this study has been deposited in the NCBI’s Sequence Read Archive (SRA) database (PRJNA1021754).

### Fluorescence in situ hybridization (FISH)

Cy3 labeled *miR-802* probe was designed and synthesized by GenePharma (Shanghai, China). The frozen sections of adipose tissue from obese patients, normal persons or obese mice were fixed with 4% formaldehyde at room temperature for 10 min. The probe was hybridized at 37℃ for 16 h. DAPI was added at 1:5000 for 15 min after washing with probe detergent. Images were obtained with confocal laser scanning microscope (CLSM, LSM800, Zeiss, Germany) and processed using the ZEN imaging software.

### Plasmid and shRNA construction

The coding sequences for *Traf3* (NM_001286122.1), *p65* (NM_ 009045.5), *RelB* (001290457.2), *Srebp1* (001313979.1) were amplified by PCR from full-length cDNA of mice, and then cloned in pcDNA 3.1 (+) vector (Addgene, Watertown, MA, USA). All plasmids were confirmed to be correct by sequencing. The primer sequences for PCR are listed in Supplementary Table 3.

The shRNA of *Traf3*, *Srebp1*, *p65*, *RelB* were constructed in plvx-shRNA2 lentivirus vector (Takara). The plvx-shRNA2 lentivirus vector was digested with *EcoR I* and *BamH I*. The shRNA primer sequences are listed in Supplementary Table 4.

### Luciferase assay

*miR-802* mimics/*miR-802* inhibitor (*anti-miR-802*)/miRNA NC (NC)/miRNA inhibitor NC was purchased from GenePharma (Shanghai, China). The construction of *Traf3* (both wild type and mutants) was achieved by digestion of pmir-PGLO vector (Addgene, Watertown, MA, USA) with double restriction enzymes (*Xhol I* and *Xbal I*), followed by ligation of sequences encoding the corresponding 3’UTR of the target genes. Sequences of the synthetic oligonucleotides encoding the 3’UTR of the target genes and their mutants are listed in Supplementary Table 3. 3T3-L1 cells were transfected with one of the above-mentioned plasmids using Lipofectamine 2000 (Invitrogen), according to the manufacturer’s instructions. At 48 h after transfection, the cells were lysed and the luciferase activity was assayed with a dual-luciferase reporter assay kit (Vazyme, Nanjing, China). Data are presented as the ratio of Renilla luciferase activity to firefly luciferase activity.

### Flow cytometric analysis of macrophage polarization

SVF was resuspended in 1 ml of Live/Dead Fixable Dead Cell stain (Molecular Probes) and incubated on ice for 30 min. Afterward, the cells were washed once with FACS buffer (1% BSA in 1×PBS), followed by staining with different antibodies. For flow cytometry analysis of macrophages, 1×10^6^ freshly isolated cells were triple stained with CD11b-Apc (#101211, Biolegend; 1:100), Zombie NIR^TM^ Fixable Viability Kit (#423105, Biolegend; 1:200) and F4/80-PE (#157304, Biolegend; 1:100), or stained with F4/80-FITC (#123108, Biolegend; 1:100), Cd206-APC (#141708, Biolegend;1:100), Cd86-PE (#105007, Biolegend; 1:100), or their isotype controls (Biolegend) on ice for 30 min in the dark. After staining, the cells were fixed with 2% (w/v) paraformaldehyde and stored at 4°C before analysis with FACS Celesta Cell Analyzer (BD Biosciences). Data were analyzed using FlowJo software version X.0.7 (Tree Star, Inc.).

### Mouse metabolic studies

After 12 h fasting treatment, mice fasting blood glucose (FBG) levels were examined via using a glucometer (OMRON, Japan) and fasting serum insulin (FINS) levels were tested by insulin ELISA kit (Crystal Chem, USA). And the homeostatic model assessment indices of insulin resistance (HOMA-IR) was calculated with the equation (FBG (mmol/l) ×FINS (mIU/l))/22.5. To perform the glucose tolerance tests, 2 g/kg glucose (Sigma-Aldrich, StLouis, MO, USA) was intraperitoneal (i.p.) injected into mice, whereas 0.75 U/kg insulin (Novolin R, Novo Nordisk, Bagsvaerd, Denmark) was i.p. injected into mice for insulin tolerance tests. Blood glucose levels were examined at 0, 15, 30, 60, 90 and 120 min after glucose or insulin injection and serum sample was collected from eye canthus blood at 0, 5, 15, and 30 min after glucose injection. Insulin level was evaluated using mice insulin ELISA kit (Crystal Chem, USA), according to the manufacturer’s instructions. Subtracting the baseline area, by subtracting the starting glucose value from the value at each time point, generates the area of the curve (AOC)(51).

### Body Composition

The changes in body composition were assessed as we have previously describe(52). In brief, mice were anesthetized with 2% isoflurane by volume in a box and fixed on an MRI platform (Bruker BioSpec 7T/20 USR). Anesthesia was also maintained with isoflurane of 1% by volume. After turning on the instrument, the mice were scanned layer by layer according to the cross-section of their internal adipose tissue content, using ImageJ software analysis and statistics of the lipid distribution in mice.

### RNA isolation and qRT-PCR analysis

Total RNA from adipose tissues or its fractions or 3T3-L1 cells was extracted with TRIzol reagent (Invitrogen). For mRNA expression analysis, 500 ng of total RNA was used for synthesis of cDNA using PrimeScriptTM RT reagent Kit (Takara, Tokyo, Japan). For miRNA expression analysis, 150 ng total RNA was reverse-transcription into cDNA using miRNA-specific primers supplied with TaqMan MicroRNA Reverse Transcription kit. The quantitative real-time PCR was performed using the LightCycle 480 (Roche). The relative level of gene expressions were calculated by the 2^-ΔΔCT^ method, after normalization with the abundance of *18S* rRNA or *U6*. For miR-802-5p and *U6*, TaqMan probes (Ambion) were used to confirm our results. The sequences of genes were listed in Supplementary Table 5.

### Western blot analysis

Proteins were extracted from tissues or cells in radioimmunoprecipitation assay (RIPA) buffer (Beyotime) containing a complete protease inhibitor cocktail (Roche), resolved by SDSPAGE, transferred onto polyvinlidene fluoride (PVDF) membranes (Bio-Rad), and then probed with primary antibodies against TRAF3 (#ab36988), NIK (#ab314146) were from Abcam, NF-κB2 p100/p52 (#4882), NF-κB p65 (#8242) were from CST, and RelB (#A23389), P52 (#ab125611), IKK-α (#A2062), phospho-IKK-α (#AP0546), IKBα (#A19714), phosho-IKBα (#AP0614), iNOS (#A14031), phosho-p65 (#AP0123), Tubulin (#AC008), β-Actin (#AC038) and Histone H3 (#A2348) were from ABclonal. The protein bands were visualized with enhanced chemiluminescence reagents (GE Healthcare) and quantified by using the ImageJ software.

### RNA immunoprecipitation (RIP)

RNA immunoprecipitation was performed using an EZMagna RIP Kit (Millipore, Billerica, MA, USA) following the manufacturer’s protocol. 3T3-L1 cells transfected with *miR-802* or *oe-Traf3* were lysed in complete RIP lysis buffer, and then, 100 μl of whole cell extract was incubated with RIP buffer containing magnetic beads conjugated with anti-Ago2 (#ab186733, Abcam) antibody or negative control normal mouse IgG (#ab172730, Abcam). Furthermore, purified RNA was subjected to qRT-PCR analysis to demonstrate the presence of the binding targets using the respective primers. The primer sequences are listed in supplementary table 5.

### Chromatin immunoprecipitation assay (ChIP)

ChIP experiments were strictly performed according to the manual for the ChIP Assay Kit (#17-10086, Millipore) and the manufacturer’s protocol. 3T3-L1 cells transfected with *miR-802* mimics, *miR-802* inhibitor, *oe-Traf3*, or *miR-802 & oe-Traf3* were fixed with 37% formaldehyde for 10 min, followed by 30 rounds of sonication, each for 3 s, to fragment the chromatin. The chromatin was incubated with NF-κB p65 antibody (#8242, CST) at 4°C overnight and then immunoprecipitated with Proteinase K (Millipore). Purified DNA was amplified by PCR using primer pairs that spanned the predicted *p65* binding sites on the *Srebp1* promoter. The primer sequences are listed in Supplementary Table 5.

### Agarose-oligonucleotide pull-down assay

The oligonucleotides for the mouse *Srebp1*a promoter and their complementary strands were synthesized by GenePharma (Shanghai, China) and biotinylated using a Pierce™ Biotin 3’ End DNA Labeling Kit (Cat. #89818; Thermo Scientific). These oligonucleotides were annealed to form double-stranded oligonucleotides, which were then incubated with streptavidin-conjugated agarose beads at 4°C for 60 min and washed twice with IP lysis buffer. Next, the nuclear extract (50 μg each) in 200 μl IP lysis buffer was pre-cleared with agarose beads at 4°C for 90 min to reduce any nonspecific binding and then incubated with oligo/streptavidin-conjugated beads at 4°C overnight. The mixtures were washed three times with IP lysis buffer via centrifugation the following day, and the affinity-purified proteins were eluted by boiling in SDS sample buffer for 10 min. Samples were then subjected to analysis by western blot. Primer sequences are listed in Supplementary Table 5.

### Histological and immunochemical analysis

The white adipose tissue was fixed in 4 % formalin solution at 4°C for 24 hours, embedded in paraffin, and sectioned at 5 μm. Deparaffinized and rehydrated sections were stained with haemotoxylin and eosin (Sigma), or with reagents for Sirius red staining, or immunohistological staining of HCS LipidTOX™ red neutral stain (#H34467, Invitrogen) and F4/80 (#GB113373, Servicebio, china). The slides were analyzed using a confocal laser scanning microscope (CLSM, Carl Zeiss LSM800) at ×20 magnification.

### Transmission electron microscopy

For transmission electron microscopy, mouse epiWAT was dissected, sliced into small fragments of 1–2 mm each, and then fixed in 5% glutaraldehyde for 2 days. Specimens were post-fixed in 1% osmium tetroxide. After staining with 2% aqueous uranyl acetate for 2 h, the samples were dehydrated in a series of ethanol up to 100% and embedded in epoxy resin. Ultrathin sections were cut with an EM UC7 ultramicrotome (Leica) and poststained with lead nitrate. Ultrathin sections were mounted in formvar-coated nickel grids and observed under an FEI Tecnai G2 Electron Microscope (FEI Tecnai G2).

### EdU labeling

Cell proliferation was detected with BeyoClick™ EdU Alexa Fluor 488 Imaging Kit (Beyotine, China). Briefly, 1×10^5^ macrophage cells or 1×10^4^ RAW264.7 cells were plated on twenty-four well plates. After co-cultured with 3T3-L1 cells or SVF cells, cells were gently washed twice with PBS, and further incubated with 10 μM EdU for 4 h. Treated cells were fixed in 4% paraformaldehyde solution at room temperature for 15 min and EdU detection was carried out according to manufacturer’s instructions.

### Sample size and replication

Sample size varied between experiments, depending on the number of mice allocated for each experiment. The minimum sample size was three.

### Data inclusion/exclusion criteria

All patients enrolled in this study were obese (BMI > 25). The negative controls were normal-weight individuals (20 ≤ BMI ≤ 25). Data or samples were not excluded from analysis for other reasons.

### Randomization

Mice used for the experiments were randomly selected and randomly assigned to experimental groups. There was no requirement for randomization of cell selection.

### Blinding

During experimentation and data acquisition, blinding was not applied to ensure tractability.

### Statistical analysis

All in vivo experiments represent individual mice as biological replicates. The exact values of *n* are reported in figure legends. Data are presented as mean ± SEM. Comparisons were performed using the Student’s *t* test between two groups or ANOVA in multiple groups. Dunn’s multiple comparisons for one-way ANOVA and Fisher’s least significant difference (LSD) for two-way ANOVA were used. The level of significance was set at *\*p* < 0.05, *\*\*p* < 0.01, *\*\*\*p* < 0.001. Graphpad prism 8 (GraphPad, San Diego, CA, USA) was used for all calculation.

## Acknowledgements

This work was supported by the National Natural Science Foundation of China: Grant No. 82373925, 82070801 (To L.J.), 82100858, 82370804 (To FF.Z.), 82073227 (To Y.P). Supported by Natural Science Foundation of Jiangsu Province, BK20221520(To L.J.), BK20200569 (To FF.Z.). Supported by grants from the ‘111’ project, B16046 (To L.J.). Supported by the Priority Academic Program Development of Jiangsu Higher Education Institutions, PAPD (To L.J.), 2632023TD03 (to FF Z). Supported by China Postdoctoral Science Foundation, 2022T150726, Supported by the Fundamental Research Funds for the Central Universities 2020M671661 (To FF.Z.) and 2632023GR07 (To Y. Y). Supported by Jiangsu Province Research Founding for Postdoctoral, 1412000016 (To FF.Z.). We would like to thank Xiaonan Ma for providing technical assistance of Carl Zeiss LSM 800 on the Public Experimental Platform of China Pharmaceutical University. We thank Yumeng Shen (Public Platform of State Key Laboratory of Natural Medicines, China Pharmaceutical University) for her assistance with flow analysis. We thank LetPub (www.letpub.com) for its linguistic assistance during the preparation of this manuscript.

## Author contributions

Y. Y, B. H and YM. Q performed the experiments; DW. W performed partial experiments on animals; YN. J and LM. S collected all the human samples; Y. P, YF. Z, YM. S and WJ. Hu analyzed data; FF. Z and L.J designed the project, FF. Z, L.J and ZY. C interpreted the data and wrote the manuscript.

---

**Figure 5-source data 1:** The original files of the full raw unedited blots of TRAF3 and β-Actin in human subcutaneous adipose tissues from obese and normal individuals (*n*_normal_=4 and *n*_obesity&IR_=9).

**Figure 5-source data 2:** The original files of the full raw unedited blots of TRAF3 and β-Actin in the epiWAT of HFD mice (*n*=3).

**Figure 5-source data 3:** The original files of the full raw unedited blots of TRAF3 and β-Actin in the epiWAT of *miR-802* KI mice (*n*=3).

**Figure 5-source data 4:** The original files of the full raw unedited blots of TRAF3 and β-Actin in 3T3-L1 cells transfected with *miR-802* mimics or *miR-802* inhibitor.

**Figure 5-source data 5**: The original files of the full raw unedited blots of TRAF3 and β-Actin in the epiWAT of control, *miR-802* KI, *Traf3^eWAT^ ^KI^*, and *miR-802* KI & *Traf3^eWAT^ ^KI^* mice (*n*=3).

**Figure 6-source data 1:** The original files of the full raw unedited blots of TRAF3, NIK and β-Actin in 3T3-L1 cells transfected with *miR-802* mimics or *miR-802* inhibitor.

**Figure 6-source data 2:** The original files of the full raw unedited blots of TRAF3, NIK and β-Actin in the epiWAT of *miR-802* KI mice (*n*=3).

**Figure 6-source data 3:** The original files of the full raw unedited blots of p100/p52 and β-Actin in 3T3-L1 cells transfected with *miR-802* mimics or *miR-802* inhibitor.

**Figure 6-source data 4:** The original files of the full raw unedited blots of p100/p52 and β-Actin in the epiWAT of *miR-802* KI mice (*n*=3).

**Figure 6-source data 5**: The original files of the full raw unedited blots of P-IKK-α, IKK-α and β-Actin in 3T3-L1 cells transfected with *miR-802* mimics or *miR-802* inhibitor.

**Figure 6-source data 6:** The original files of the full raw unedited blots of P-IKK-α, IKK-α and β-Actin in the epiWAT of *miR-802* KI mice (*n*=3).

**Figure 6-source data 7:** The original files of the full raw unedited blots of p100/p52, P-IKK-α, IKK-α, NIK and β-Actin in the 3T3-L1 cells.

**Figure 6-source data 8:** The original files of the full raw unedited blots of p100/p52, P-IKK-α, IKK-α, NIK and β-Actin in the epiWAT of *miR-802* KI and *Traf3^eWAT^ ^KI^* mice (*n*=3).

**Figure 6-source data 9:** The original files of the full raw unedited blots of some major canonical NF-κB signaling targets in the epiWAT of *miR-802* KI mice(*n*=3).

**Figure 6-source data 10**: The original files of the full raw unedited blots of some major canonical NF-κB signaling targets in the epiWAT of *miR-802* KI mice and *Traf3^eWAT^ ^KI^* rescued mice (*n*=3).

**Figure 7-source data 1:** The original files of the full raw unedited blots of m-SREBP1, P-SREBP1 and β-Actin in mature 3T3-L1 cells transfected with *miR-802* mimics or *miR-802* inhibitor.

**Figure 7-source data 2:** The original files of the full raw unedited blots of m-SREBP1, P-SREBP1 and β-Actin in in the epiWAT of *miR-802* KI mice (*n*=3).

**Figure 7-source data 3:** The original files of the full raw unedited blots of m-SREBP1, P-SREBP1 and β-Actin in the 3T3-L1 cells.

**Figure 7-source data 4:** The original files of the full raw unedited blots of p65 and β- Actin in 3T3-L1 cells transfected with *miR-802* mimics or *miR-802* inhibitor.

**Figure 7-source data 5**: The original files of the full raw unedited blots of p65 and β- Actin in 3T3-L1 cells stimulated with *p65* plasmid for 48 h.

**Figure 2-figure supplement 2-source data 1**: The original files of the full raw unedited gels of *miR-802* KI mice.

**Figure 3-figure supplement 3-source data 1**: The original files of the full raw unedited gels of *miR-802* KO mice.

**Figure 5-figure supplement 5-source data 1:** The original files of the full raw unedited blots of TRAF3 and β-Actin in the epiWAT of *ob/ob* mice (*n*=3).

**Figure 5-figure supplement 5-source data 2:** The original files of the full raw unedited blots of TRAF3 and β-Actin in the epiWAT of *db/db* mice (*n*=3).

**Figure 5-figure supplement 5-source data 3:** The original files of the full raw unedited blots of TRAF3 and β-Actin in the epiWAT of *miR-802* KO mice (*n*=3).

**Figure 6-figure supplement 6-source data 1:** The original files of the full raw unedited blots of TRAF3, NIK and β-Actin in in the epiWAT of *miR-802* KO mice (*n*=3).

**Figure 6-figure supplement 6-source data 2:** The original files of the full raw unedited blots of p100/p52 and β-Actin in the epiWAT of *miR-802* KO mice (*n*=3).

**Figure 6-figure supplement 6-source data 3:** The original files of the full raw unedited blots of P-IKK-α, IKK-α and β-Actin in the epiWAT of *miR-802* KO mice (*n*=3).

**Figure 7-figure supplement 7-source data 1:** The original files of the full raw unedited gels by ChIP-PCR experiments in the 3T3-L1 cells.

**Figure 7-figure supplement 7-source data 2:** The original files of the full raw unedited gels by ChIP-PCR experiments in the epiWAT of *miR-802* KI mice (*n*=3).

## Supplemental information

**Supplementary Figure 1.**
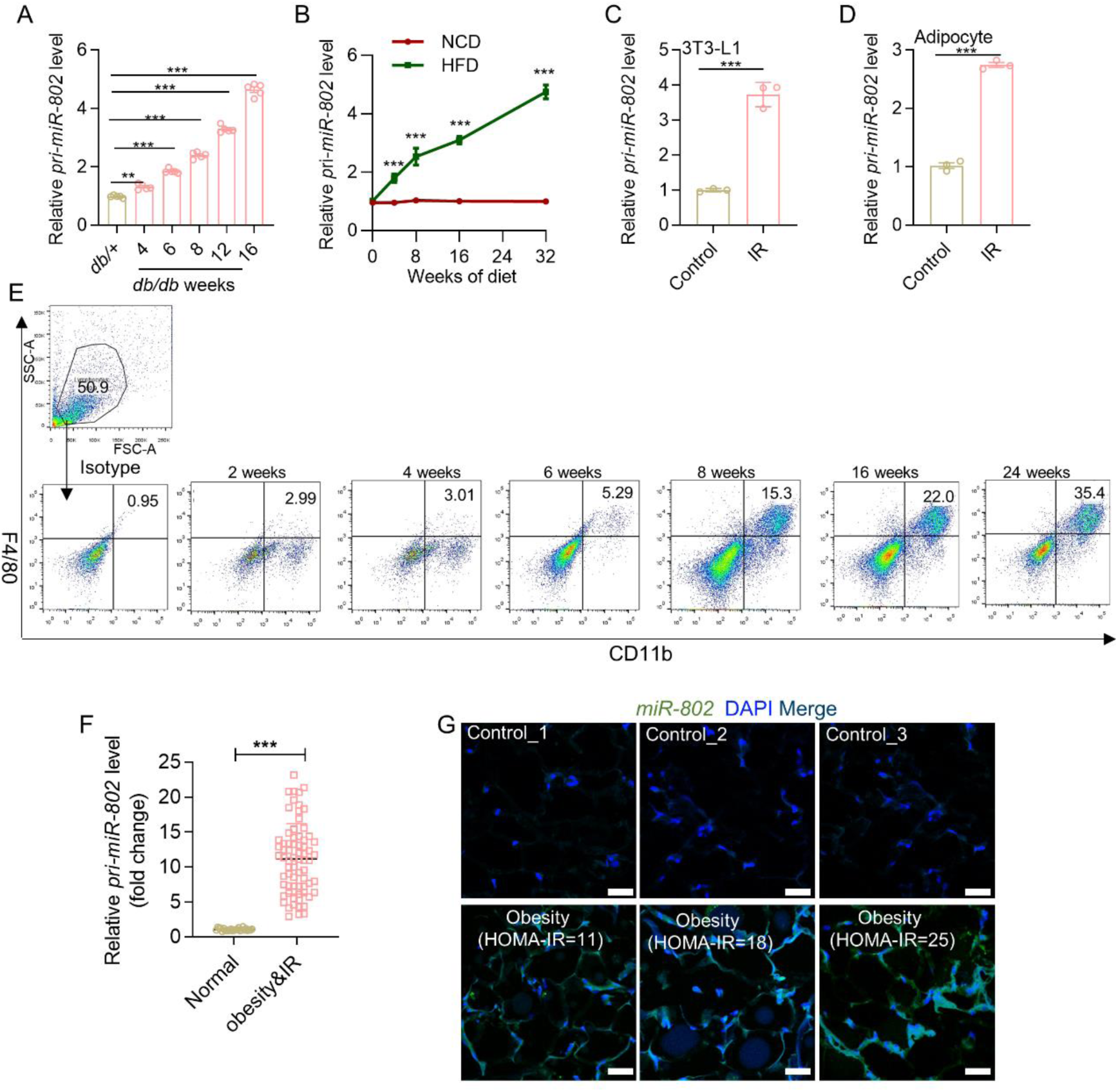
(A) The mRNA abundance of *pri-miR-802* in epiWAT of 4, 6, 8, 12, 16 weeks *db/db* mice or control mice (*n*=5). (B) The mRNA abundance of *pri-miR-802* in epiWAT of mice fed with normal chow diet (NCD) or HFD for 0, 2, 4, 8, 16, 24 and 32 weeks (*n*=5). (C-D) The insulin-resistant cell models were established in 3T3-L1 (C) and WAT SVF cells (D) by 0.5 mM palmitate, 10 μg/ml insulin and 25 mM glucose for 24 h, and qRT-PCR was performed to measure the expression levels of *pri-miR-802*. (E) Representative images of flow cytometric analysis of CD11b^+^/F4/80^+^ in the SVFs isolated from eipWAT in mice fed with HFD (*n*=5). (F) The expression levels of *pri-miR-802* in the human subcutaneous adipose tissues (*n_normal_*=25, *n_obesity & IR_*=70). (G) FISH analysis of *miR-802* in the human subcutaneous adipose tissues of obese patient or normal patient (*n*=7). The nuclei were stained with DAPI. Magnification: ×20, scale bar, 20 μm. Data represent mean ± SEM. The p-values by two-tailed unpaired Student’s *t* test (C-D, F), or two-way ANOVA (A-B) are indicated. *\*\*P* < 0.01, *\*\*\*P* < 0.001. Relative levels of *pri-miR-802* were normalized to *U6*.

**Supplementary Figure 2.**
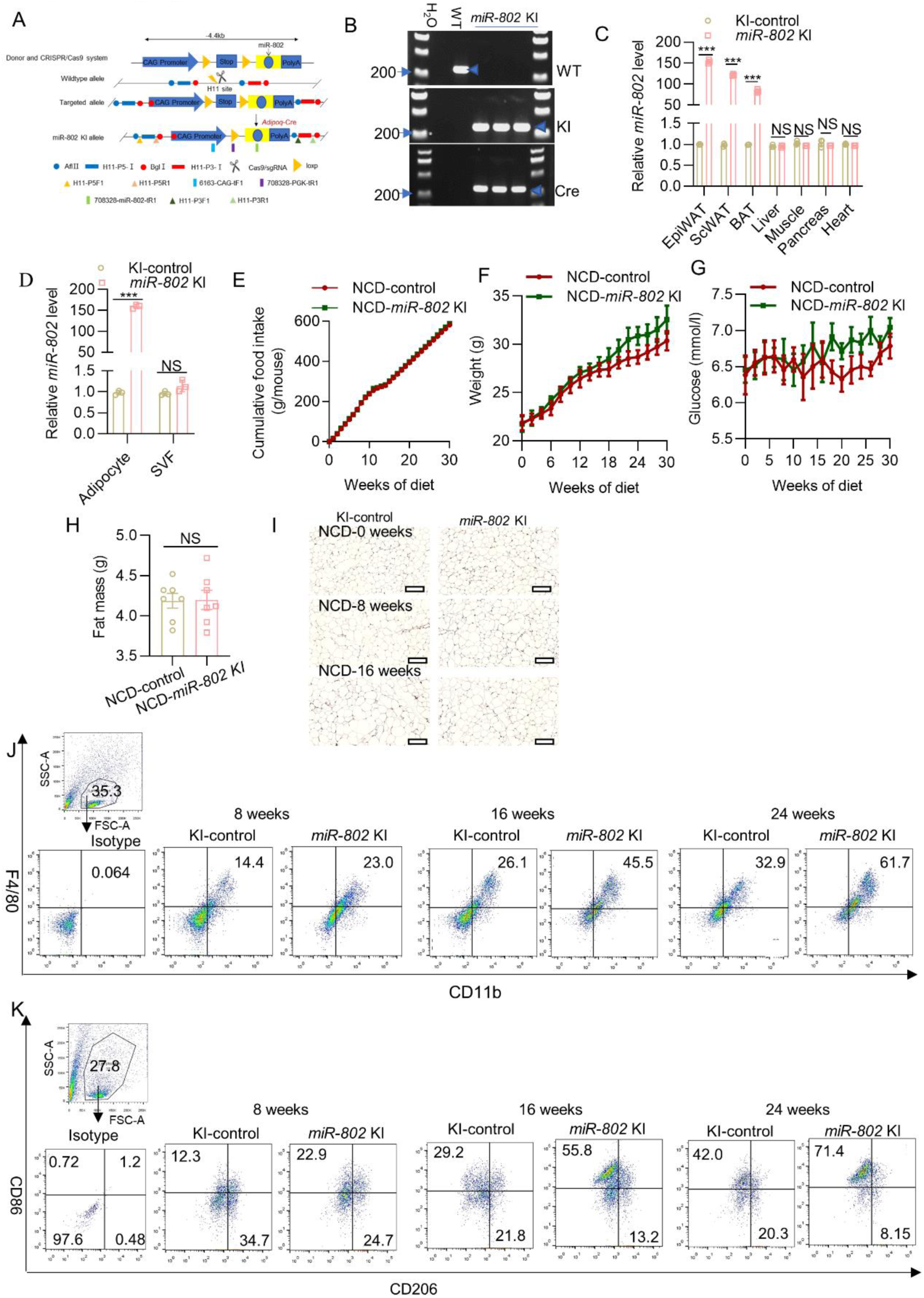
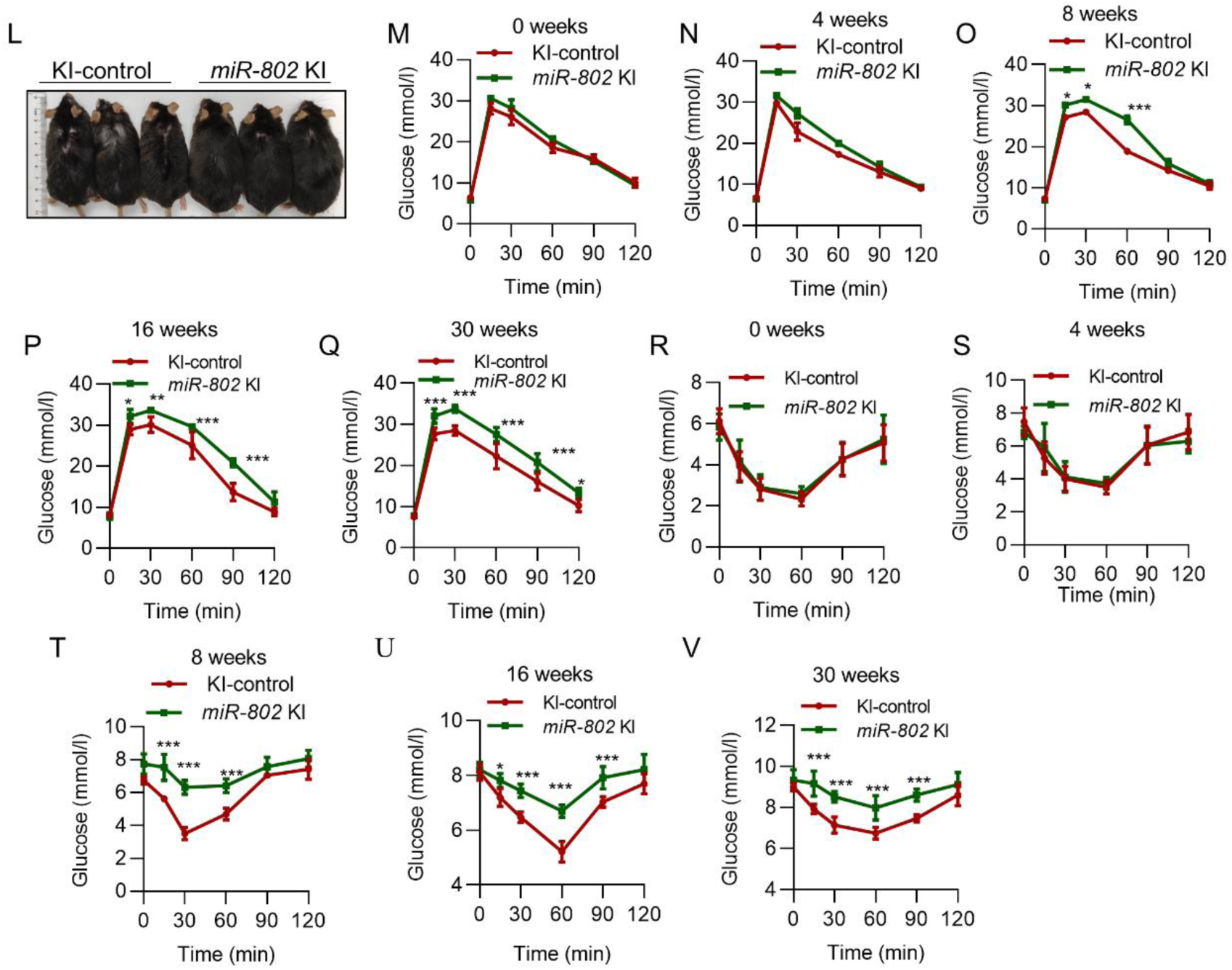
(A) Schematic diagram showing the strategy for generation of adipose-specific *miR-802* KI mice. (B) Genotypic PCR analysis showing that the adipose tissue *miR-802* WT mouse carrying homozygous *miR-802* KI allele, while KI mouse carrying both KI and Cre allele. (C) qRT-PCR analysis showing a markedly decreased expression of *miR-802* in several adipose tissues (epiWAT, scWAT, BAT), but not in liver or heart tissues (*n*=3). (D) *miR-802* mRNA levels in isolated adipocytes and SVF from epiWAT of *miR-802* KI mice (*n*=3). (E) The cumulative food intake of *miR-802* KI and control mice treated with NCD feeding (*n*=5). (F-G) Dynamic changes in body weight (F) and glucose (G) of control and *miR-802* KI mice during 30 weeks of NCD feeding (*n*=7). (H) Fat mass of whole body of control and *miR-802* KI mice of NCD feeding (*n*=7). (I) Representative images of F4/80 in epiWAT of WT or *miR-802* KI mice on NCD for 0, 8, and 16 weeks (*n*=5). Scale bar: 40 μm. (J) Representative images of flow cytometric analysis of CD11b^+^/F4/80^+^ cells in the SVFs isolated from eipWAT in control or *miR-802* KI mice fed with HFD (*n*=5). (K) Representative images of flow cytometric analysis of CD86 or CD206 in the SVFs isolate from eipWAT in control or *miR-802* KI mice fed with HFD (*n*=5). (L) Representative photos of adipose-specific *miR-802* KI mice and their WT *miR-802^ki/ki^*littermates fed with either HFD for 16 weeks (*n*=3). (M-Q) IPGTT (1.5 g/kg, K-O) and IPITT (0.75 U/kg, R-V) were performed in *miR-802 KI* mice and control mice at the 0th, 4th, 8th, 16th or 30th week of high-fat diet administration, respectively (*n*=5). Data represent mean ± SEM. Differences between groups were determined by ANOVA (C-G, M-V) or two-tailed unpaired Student’s t test (H). *\*\*\*P* < 0.001. *miR-802* abundance was normalized to U6 level.

**Supplementary Figure 3.**
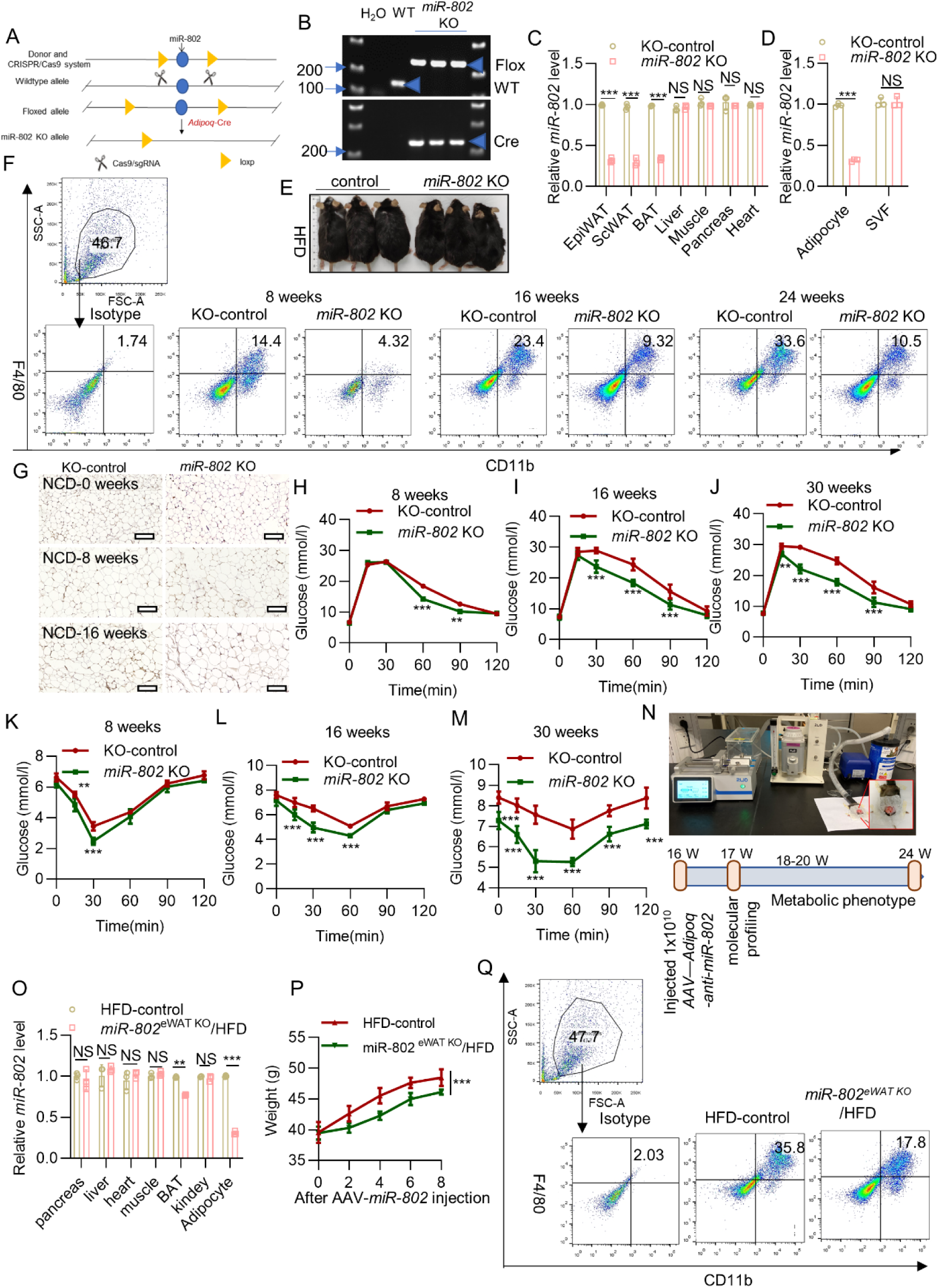
(A) Schematic diagram showing the strategy for generation of adipose-specific *miR-802* KO mice. (B) Genotypic PCR analysis showing that the adipose tissue *miR-802* WT mouse carrying homozygous *miR-802* KO allele, while *miR-802* KO mouse carrying both *miR-802* KO and Cre allele. (C) qRT-PCR analysis showing a markedly decreased expression of *miR-802* in several adipose tissues (epiWAT, scWAT, BAT), but not in liver or heart tissues (*n*=3). (D) *miR-802* mRNA levels in isolated adipocytes and SVF from epiWAT of *miR-802* KO mice (*n*=3). (E) Representative photos of adipose-specific *miR-802* KO mice and their WT *miR-802*^fl/fl^ littermates fed with either HFD for 16 weeks (*n*=3). (F) Cells isolated from SVFs of epiWAT in m*iR-802* KO and KO control mice fed with HFD for 8, 16, and 24 weeks were subjected to flow cytometry analysis for percentage of CD11b^+^/F4/80^+^ total macrophages (*n*=5). (G) Representative images of F4/80 in epiWAT of WT or *miR-802* KO mice on NCD for 0, 8, and 16 weeks (*n*=5). Scale bar: 40 μm. IPGTT (H-J) and IPITT (K-M) were performed in *miR-802* KO mice and control mice at the 8th, 16th or 30th week of high-fat diet administration, respectively (*n*=5). (N) Flowchart of the *in vivo* experiments designed for detecting adipose tissue inflammation and metabolic function via inguinal fat pad infusion of AAV-*Adipoq-anti-miR-802* (*n*=10). (O) The expression levels of *miR-802* in the different tissue of HFD-control mice or *miR-802^eWAT^ ^KO^*/HFD mice (*n*=3). (P) Dynamic changes in body weight of *miR-802^eWAT^ ^KO^*/HFD mice and control during 8 weeks of HFD feeding. (Q) Representative images of flow cytometric analysis of CD11b^+^/F4/80^+^ in the SVFs isolated from eipWAT in HFD-control or *miR-802^eWAT^ ^KO^* /HFD mice (*n*=3). Data represent mean ± SEM. Differences between groups were determined by ANOVA (C-D, H-M, O). *\*\*P* < 0.01, *\*\*\*P* < 0.001. *miR-802* abundance was normalized to *U6* level.

**Supplementary Figure 4.**
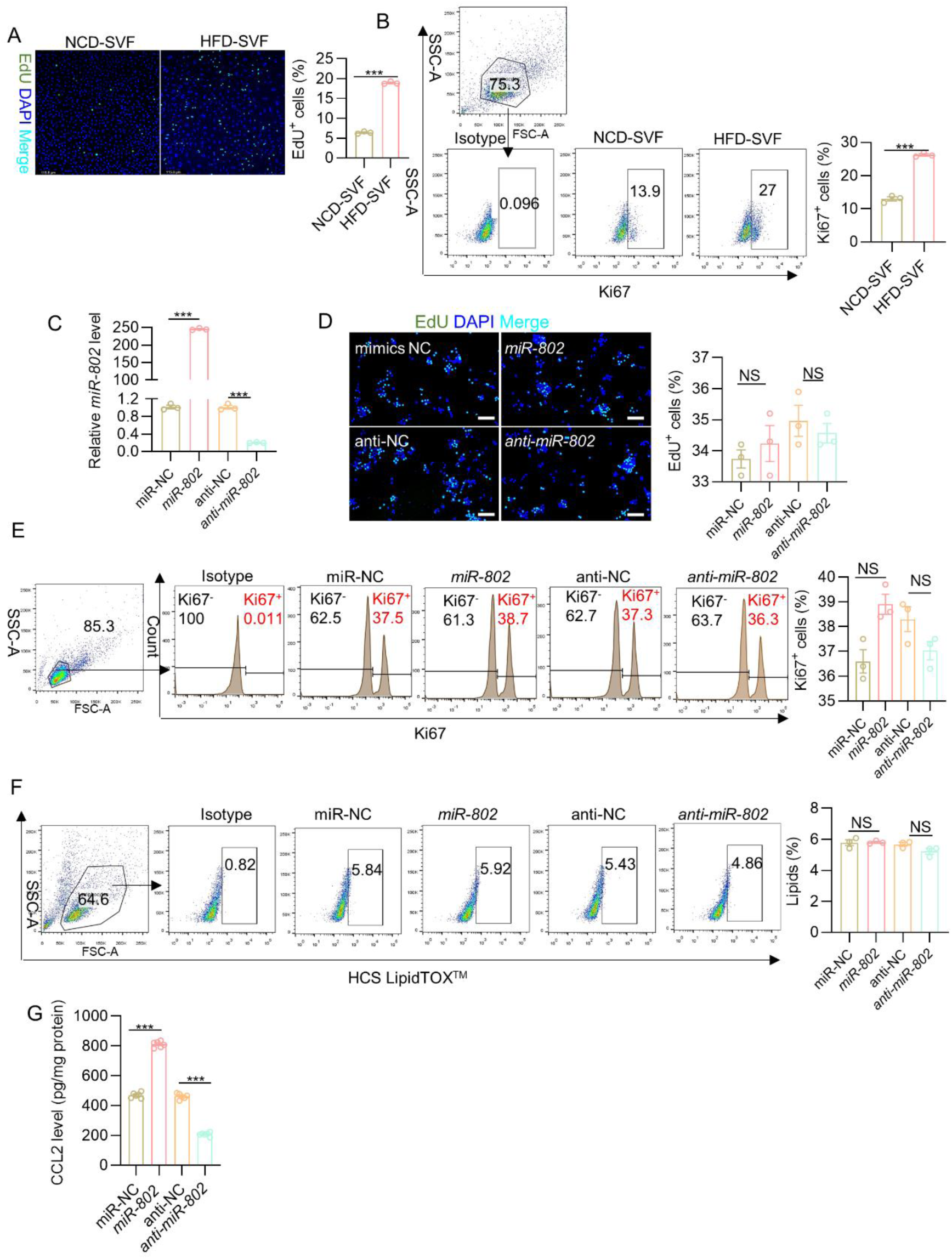
(A-B) obesity induced macrophages proliferation tested by EdU staining (A) and flow cytometry analysis (FACS, B). (C) qRT-PCR was performed to test the *miR-802* expression levels in the 3T3-L1 cells transfected with *miR-802* mimics or *miR-802* inhibitor. (D-E) EdU staining (D) and FACS analysis (E) were used to detect the proliferation of RAW264.7 cells. (F) FACS analysis of LipidTOX^TM^ in RAW 264.7 cells. (G) The CCL2 levels were determined with ELISA. Data represent mean ± SEM. Differences between groups were determined by ANOVA (C-D, G). *\*\*P* < 0.01, *\*\*\*P* < 0.001. *miR-802* abundance was normalized to *U6* level.

**Supplementary Figure 5.**
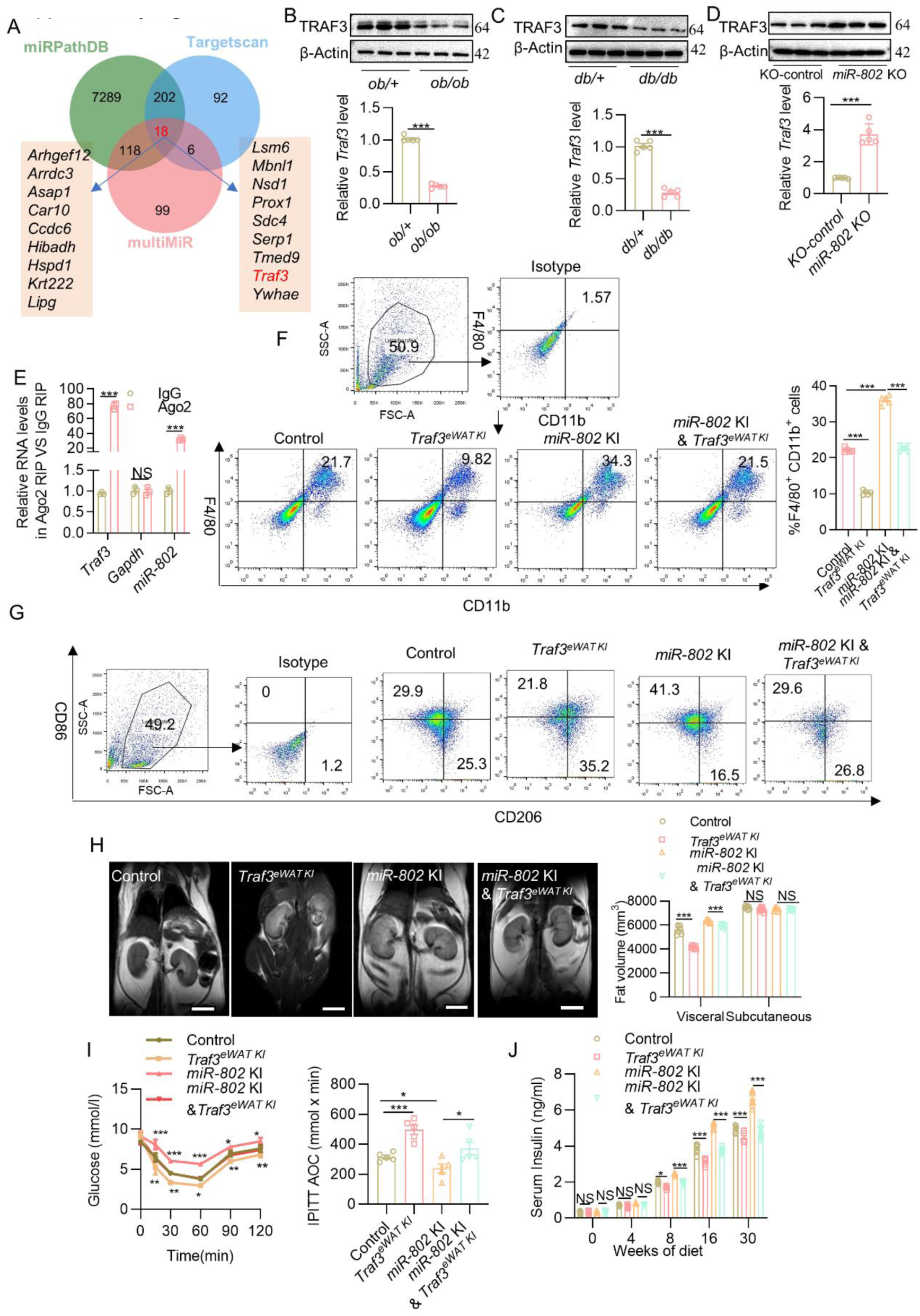
(A) miRPathDB, Targetscanand and multiMiR were used to predict the target genes of *miR-802*. (B-D) The mRNA and protein levels of TRAF3 in the epiWAT of *ob/ob* mice (B, *n*=3), *db/db* mice (C, *n*=3) or *miR-802* KO mice (D, *n*=3). (E) Anti-Ago2 RIP was performed in 3T3-L1 cells transiently overexpressing *Traf3*, followed by qRT-PCR to detect *miR-802* associated with Ago2 (nonspecific IgG served as a negative control). (F-G) Cells isolated from SVFs of epiWAT in control, *Traf3^eWAT^ ^KI^*, *miR-802* KI and *miR-802* KI & *Traf3^eWAT^ ^KI^*, mice fed with HFD 16 weeks were subjected to flow cytometry analysis for percentage of CD11b^+^/F4/80^+^ total macrophages (F, *n* =3) and M1 (CD86^+^CD206^-^) and M2 (CD206^+^CD86^-^) within the macrophage population (G, *n*=3). (H) Representative coronal section MRI images and visceral and subcutaneous adipose tissue volume of HFD-fed control, *Traf3^eWAT^ ^KI^*, *miR-802* KI and *miR-802* KI & *Traf3^eWAT^ ^KI^* mice. (I) Insulin tolerance test after mice were fed with HFD 16 weeks. (J) Serum insulin levels of control, *miR-802* KI, *Traf3^eWAT^ ^KI^* and *miR-802* KI & *Traf3^eWAT^ ^KI^* mice during 30 weeks of NCD or HFD feeding (*n*=7). Data represent mean ± SEM. Differences between groups were determined by ANOVA (E-F, I-J). *\*\*\*P* < 0.001. *miR-802* abundance was normalized to *U6* level, and other genes levels were normalized to *18S rRNA* abundance.

**Supplementary Figure 6.**
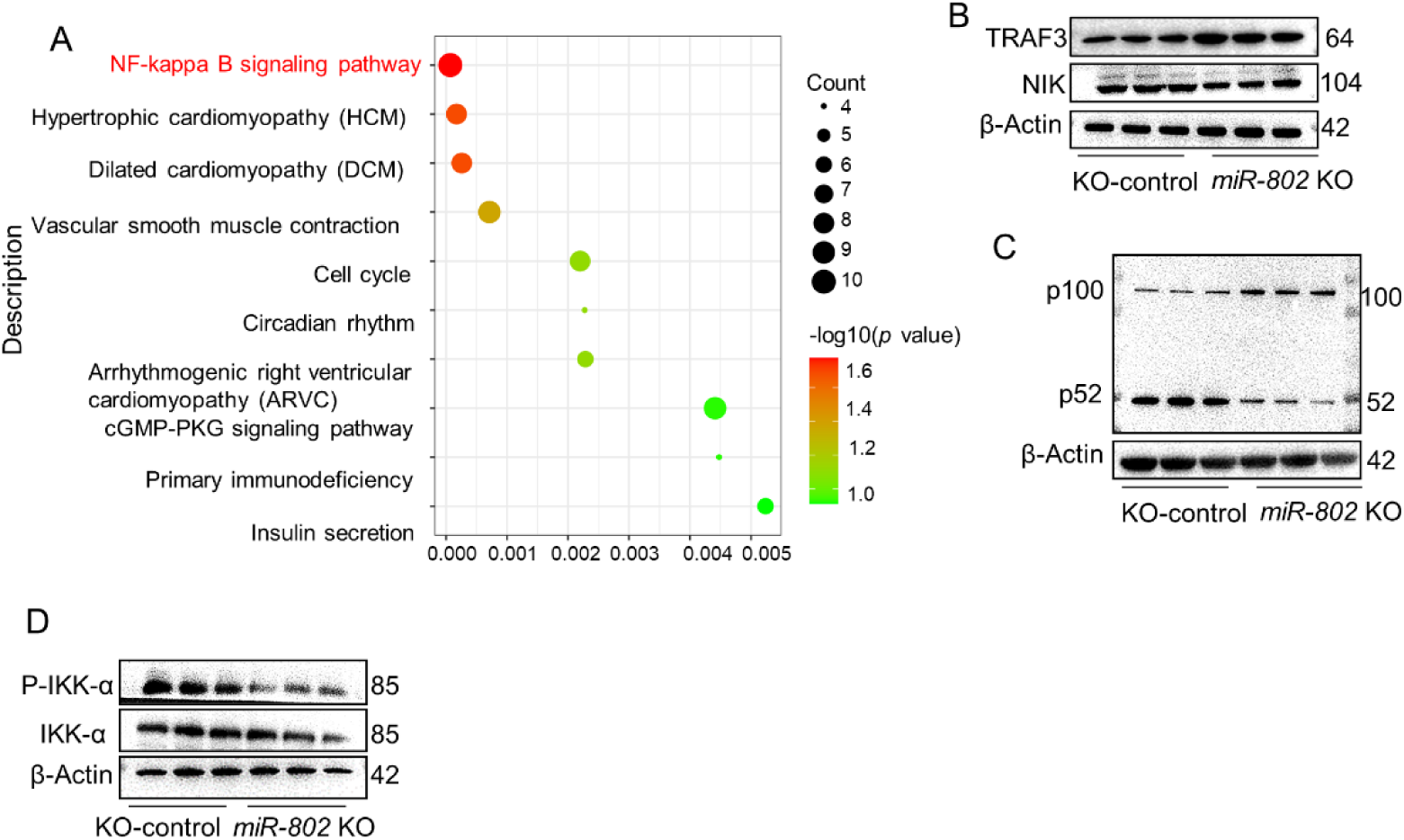
(A) GO analysis of RNA sequencing in epiWAT of *miR-802* KI mice compared to their WT *miR-802*^fl/fl^ littermates. (B) NIK protein levels in the epiWAT of *miR-802* KO mice (*n*=3). (C) P100/52 protein levels in the epiWAT of *miR-802* KO mice (*n*=3). (D) The protein levels of IKK-α and P-IKK-α in the epiWAT of *miR-802* KO mice (*n* =3). Data represent mean ± SEM.

**Supplementary Figure 7.**
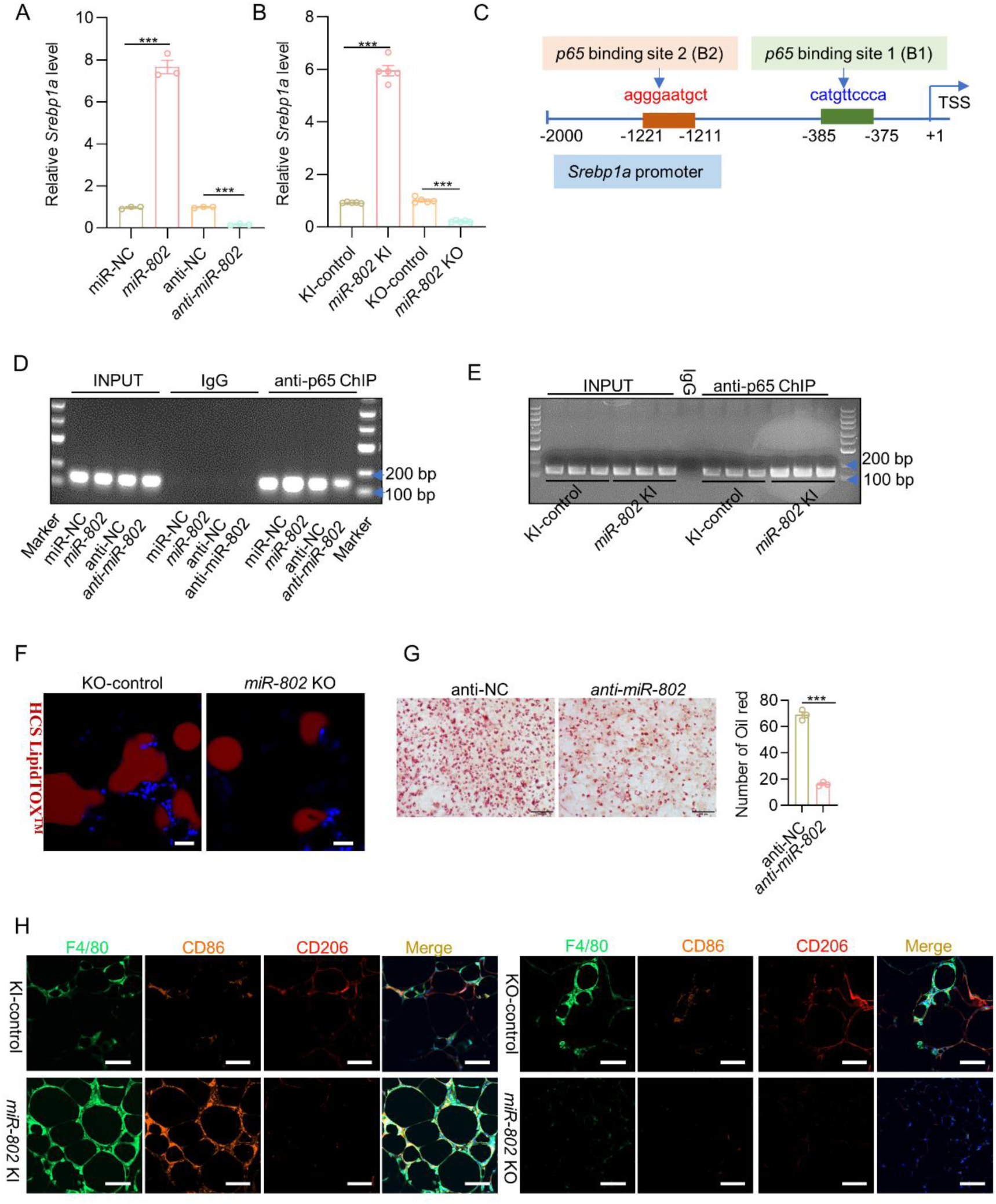
(A-B) qRT-PCR was performed to detect *Srebp1a* mRNA levels in the 3T3-L1 cells *miR-802* mimics or *miR-802* inhibitor (A) and in the epiWAT of *miR-802* KI mice (B, *n*=3). (C) The predicted binding site of *p65* on the *Srebp1* promoter. (D-E) ChIP-PCR experiments were conducted to verify that *p65* binds to the promoter of *Srebp1* in the 3T3-L1 cells (D) and in the epiWAT of *miR-802* KI mice (E, *n* = 3). (F) Representative images of immunofluorescence of lipid drop (HCS LipidTOXTM, Red) and DAPI (Blue). Scale bar: 20 μm. (G) Oil red O staining was performed to test the lipid droplet number in the 3T3-L1 cells transfected with *miR-802* inhibitor. (H) Immunohistochemical analysis was performed to test F4/80, CD86 and CD206 levels in the epiWAT of *miR-802* KI mice (*n*=3), Scale bar: 20 μm. Data represent mean ± SEM. Differences between groups were determined by ANOVA (A-B). *\*\*\*P* < 0.001. Genes levels were normalized to *18S rRNA* abundance.

**Supplementary Table 1.**
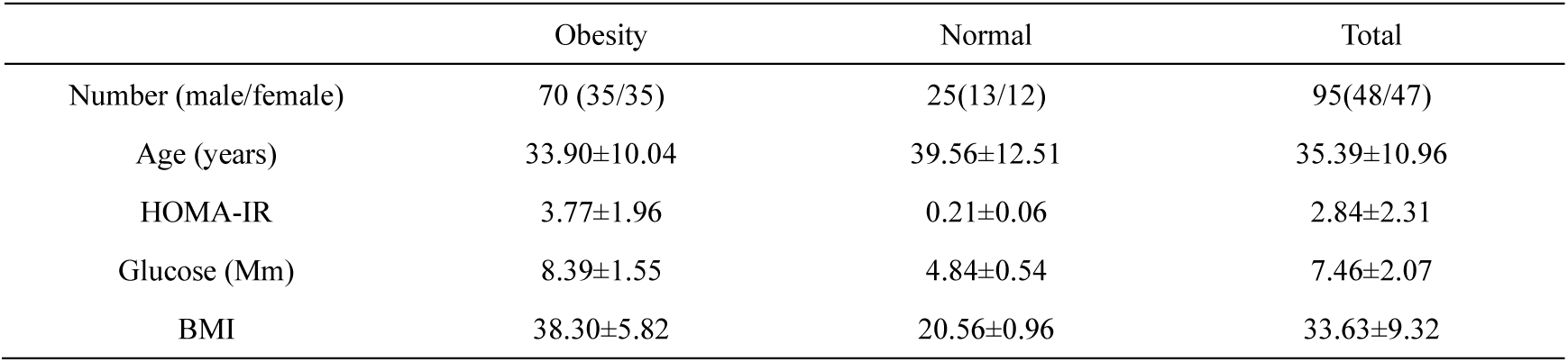
Clinical characteristics of the patients with obese patients and normal individuals.

|  | Obesity | Normal | Total |
| --- | --- | --- | --- |
| Number (male/female) | 70 (35/35) | 25(13/12) | 95(48/47) |
| Age (years) | 33.90±10.04 | 39.56±12.51 | 35.39±10.96 |
| HOMA-IR | 3.77±1.96 | 0.21±0.06 | 2.84±2.31 |
| Glucose (Mm) | 8.39±1.55 | 4.84±0.54 | 7.46±2.07 |
| BMI | 38.30±5.82 | 20.56±0.96 | 33.63±9.32 |

**Supplementary Table 2.**
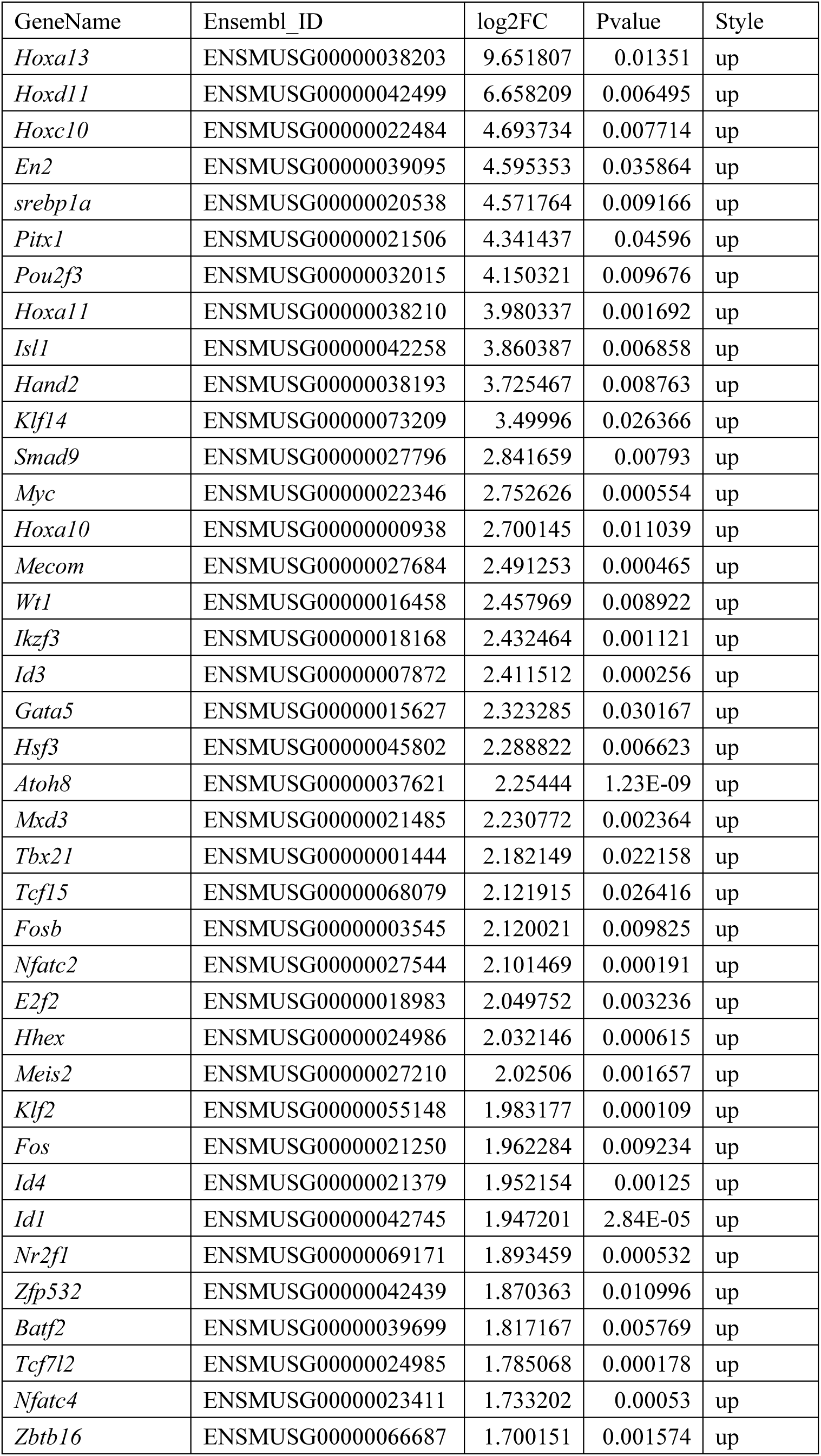
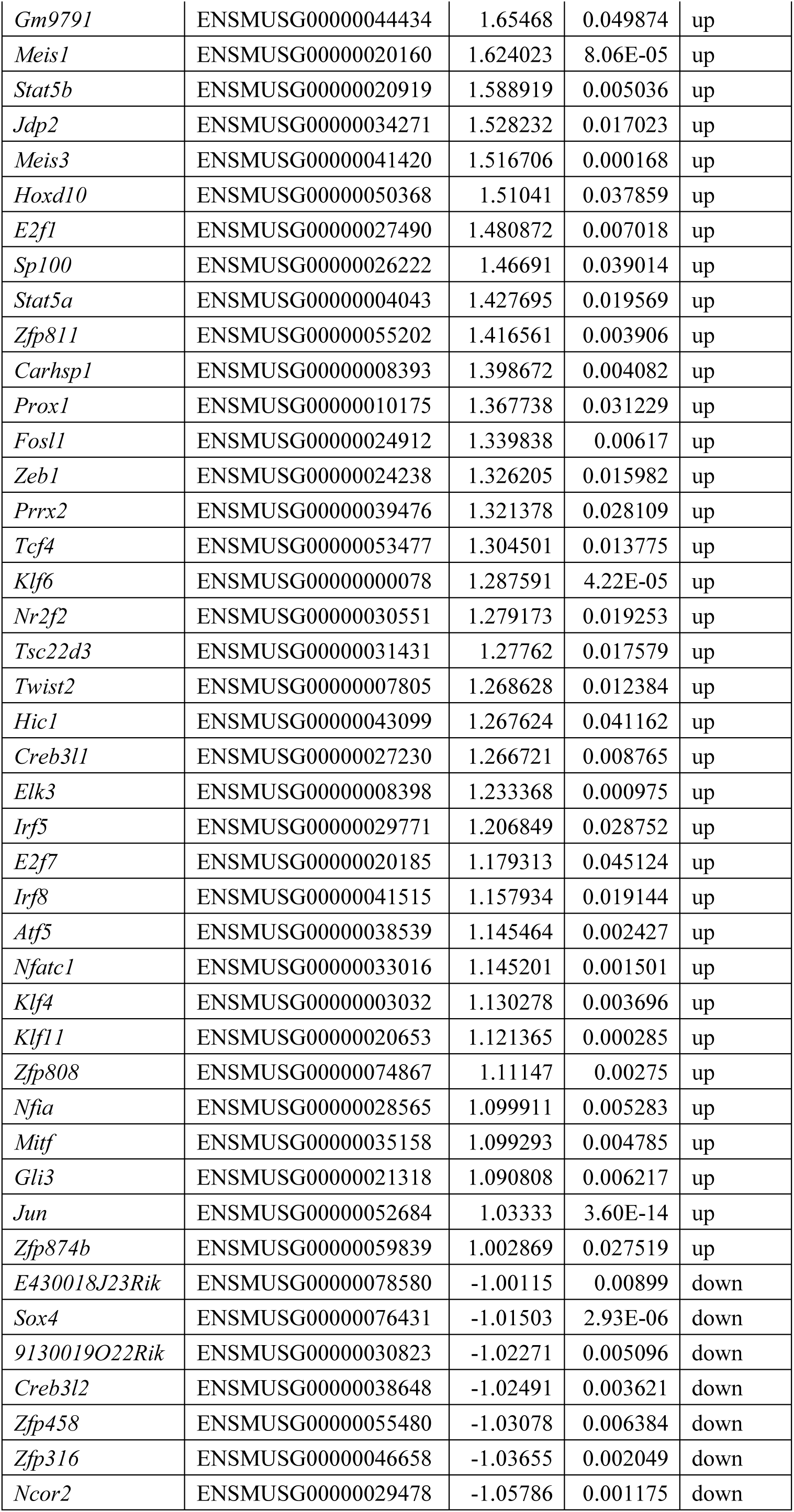
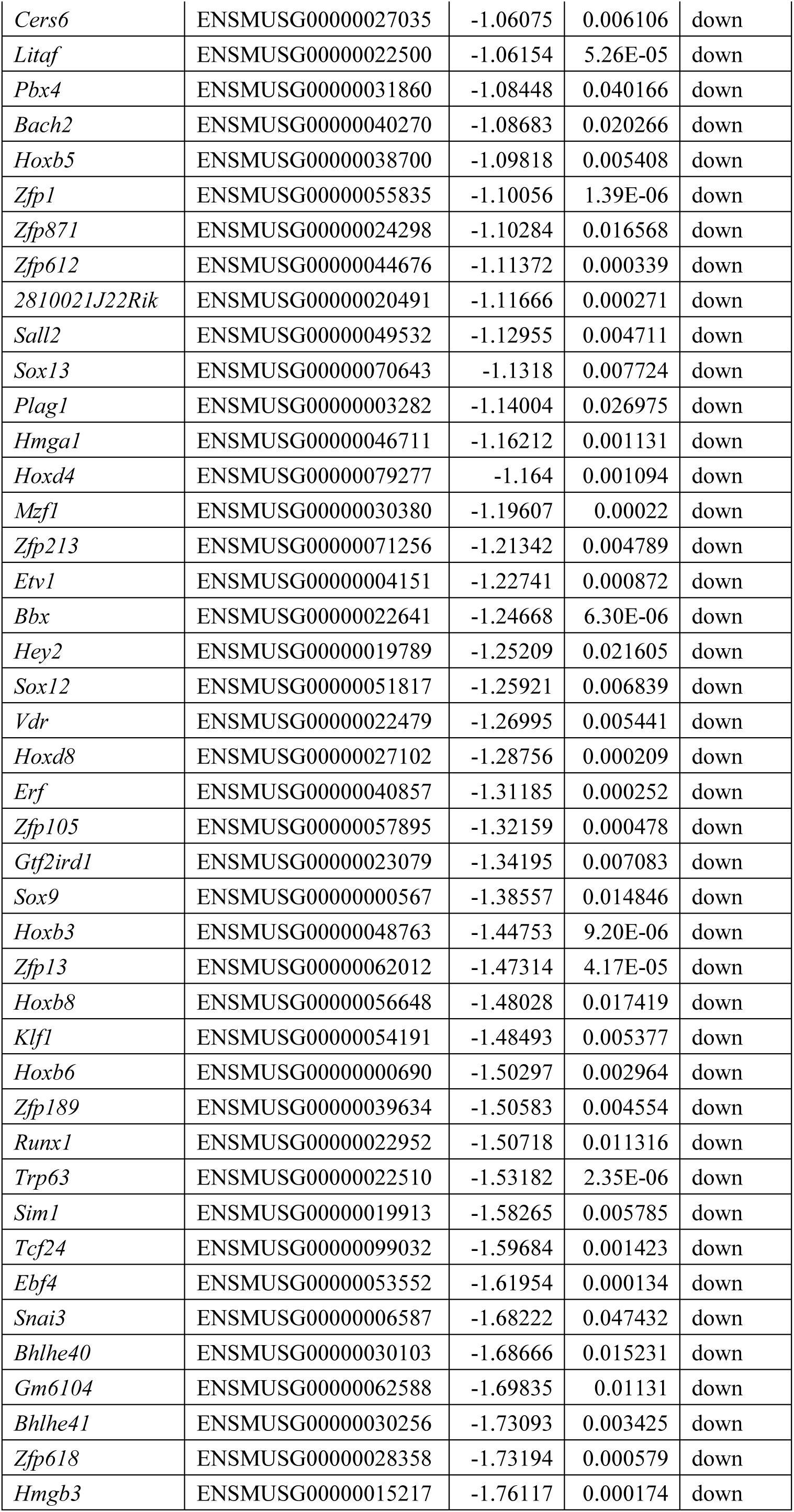
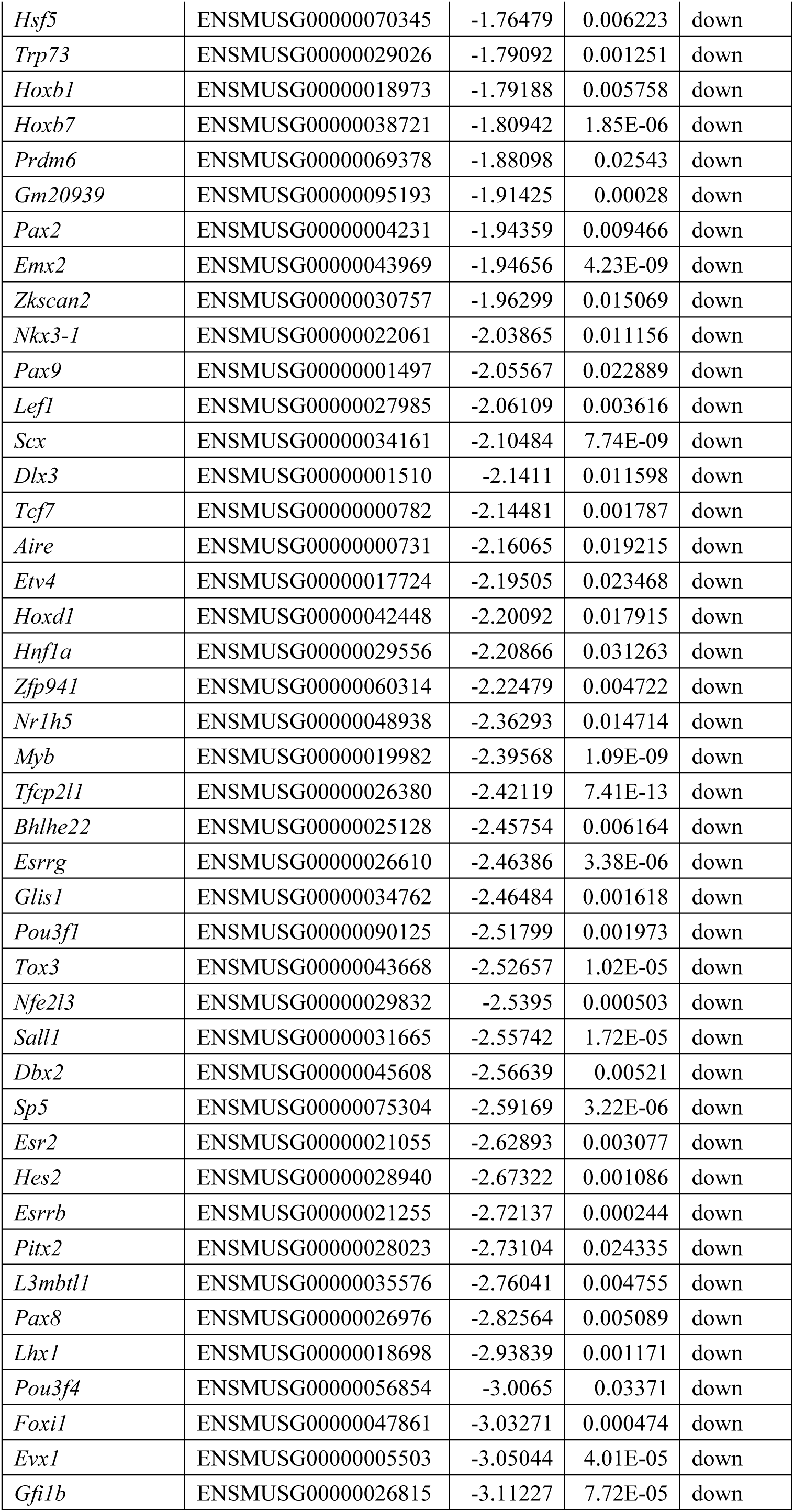
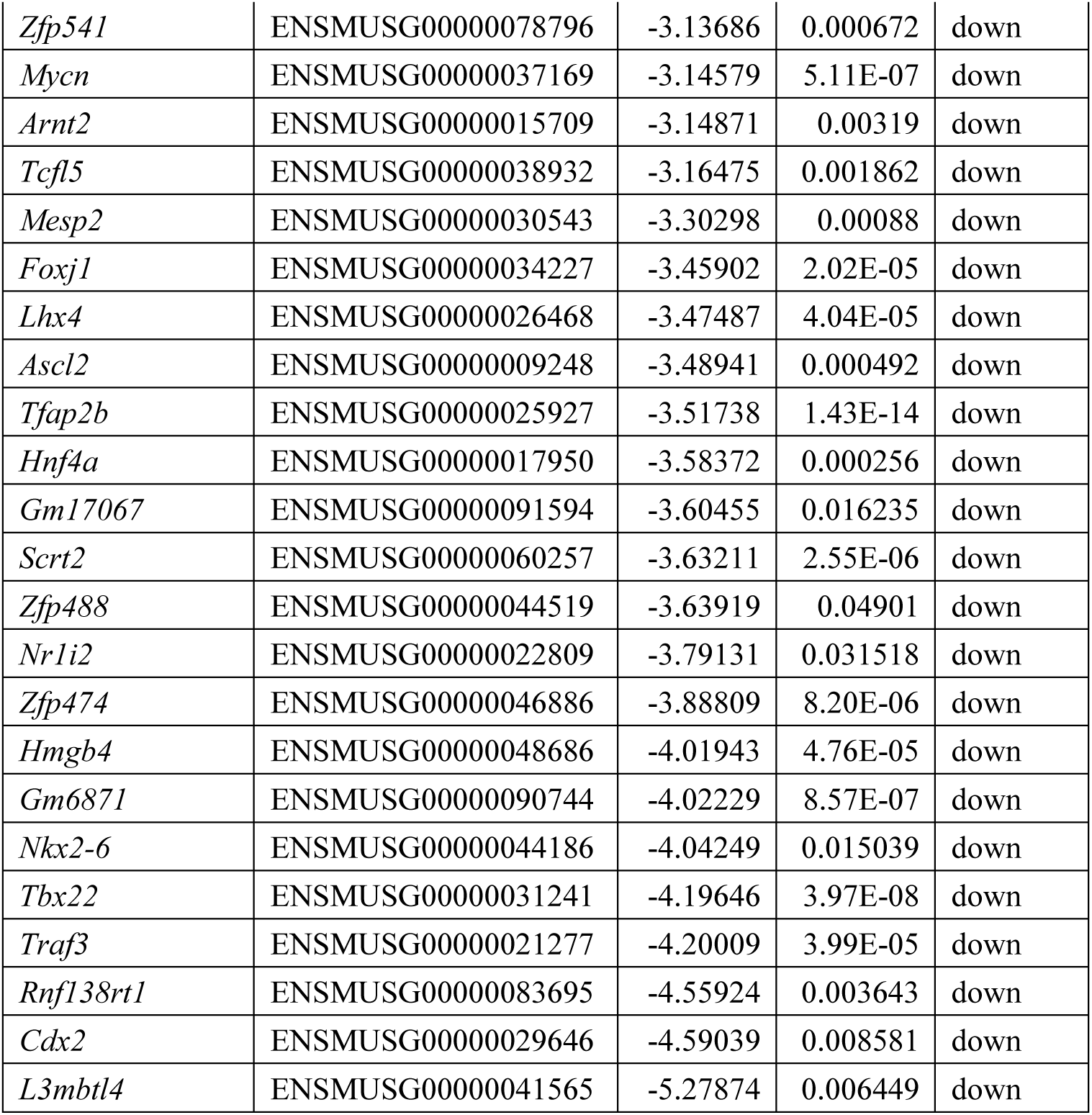
RNA islolated from epiWAT of wide type mice and *miR-802* KI mice, this table shows significantly changed mRNA (Log2 (FPKM (*miR-802* KI/WT)) ≥1).

**Supplementary Table 3.**
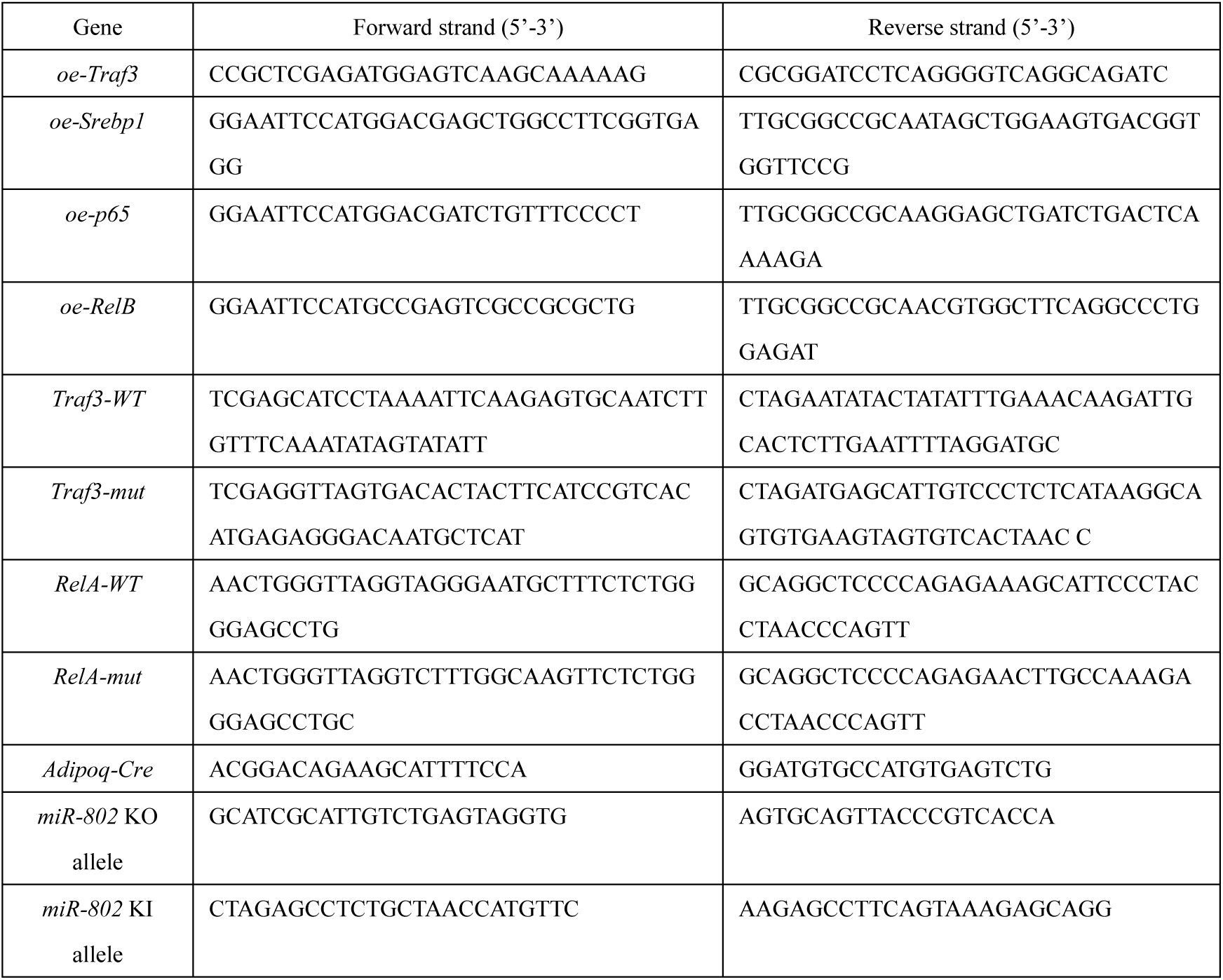
Primer sequences used for RT-PCR.

**Supplementary Table 4.**
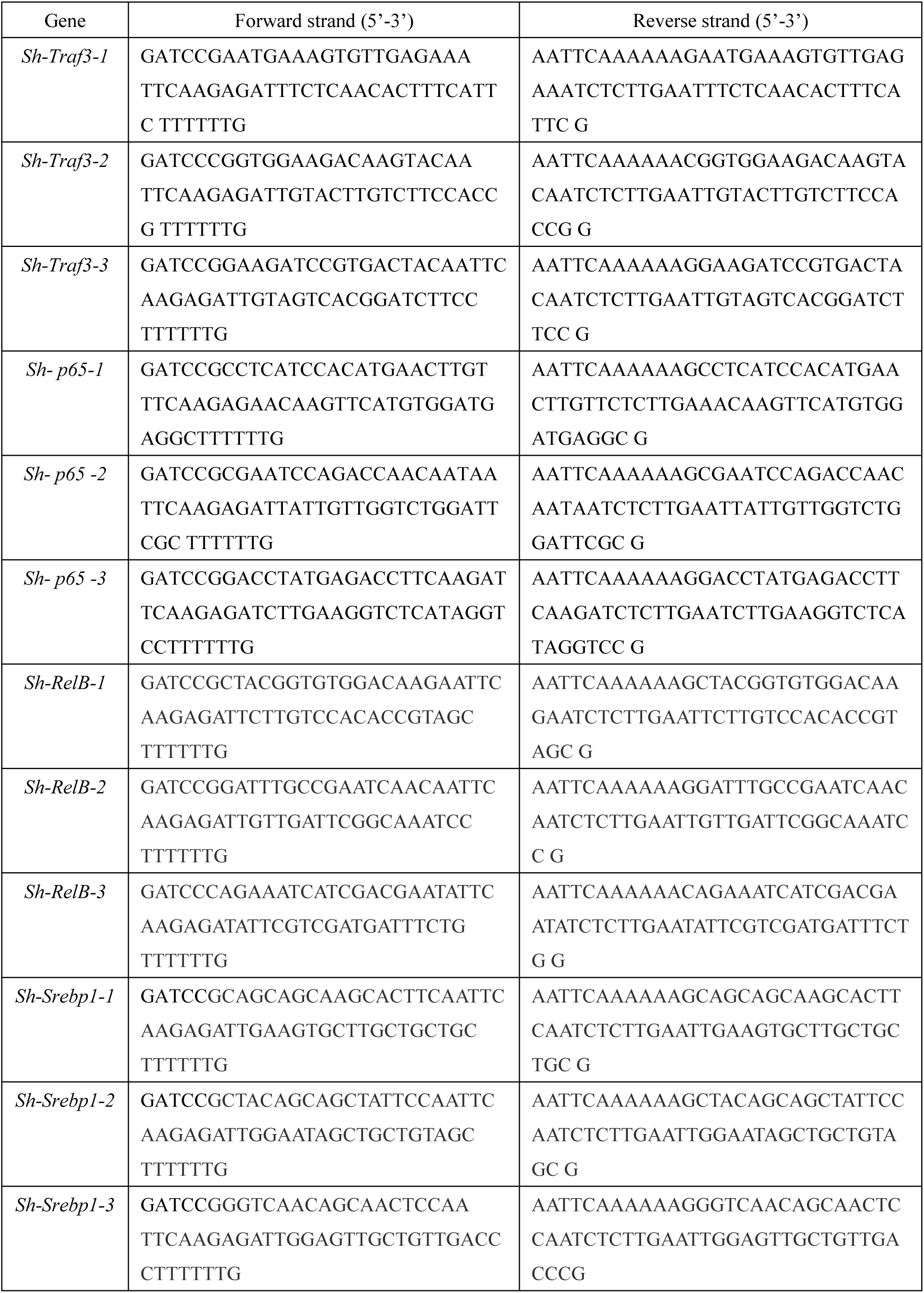
Oligo sequences used for shRNA.

**Supplementary Table 5.**
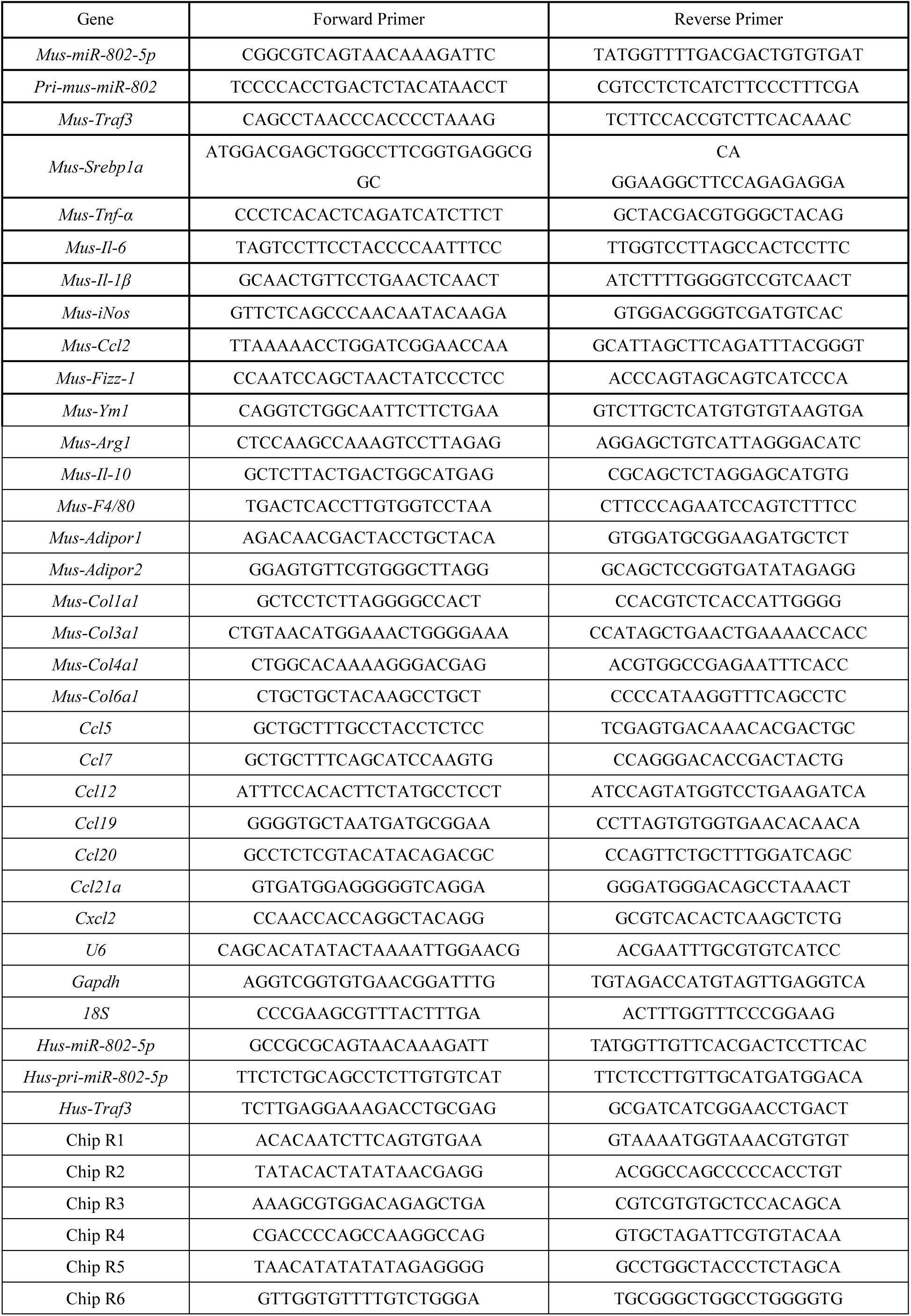
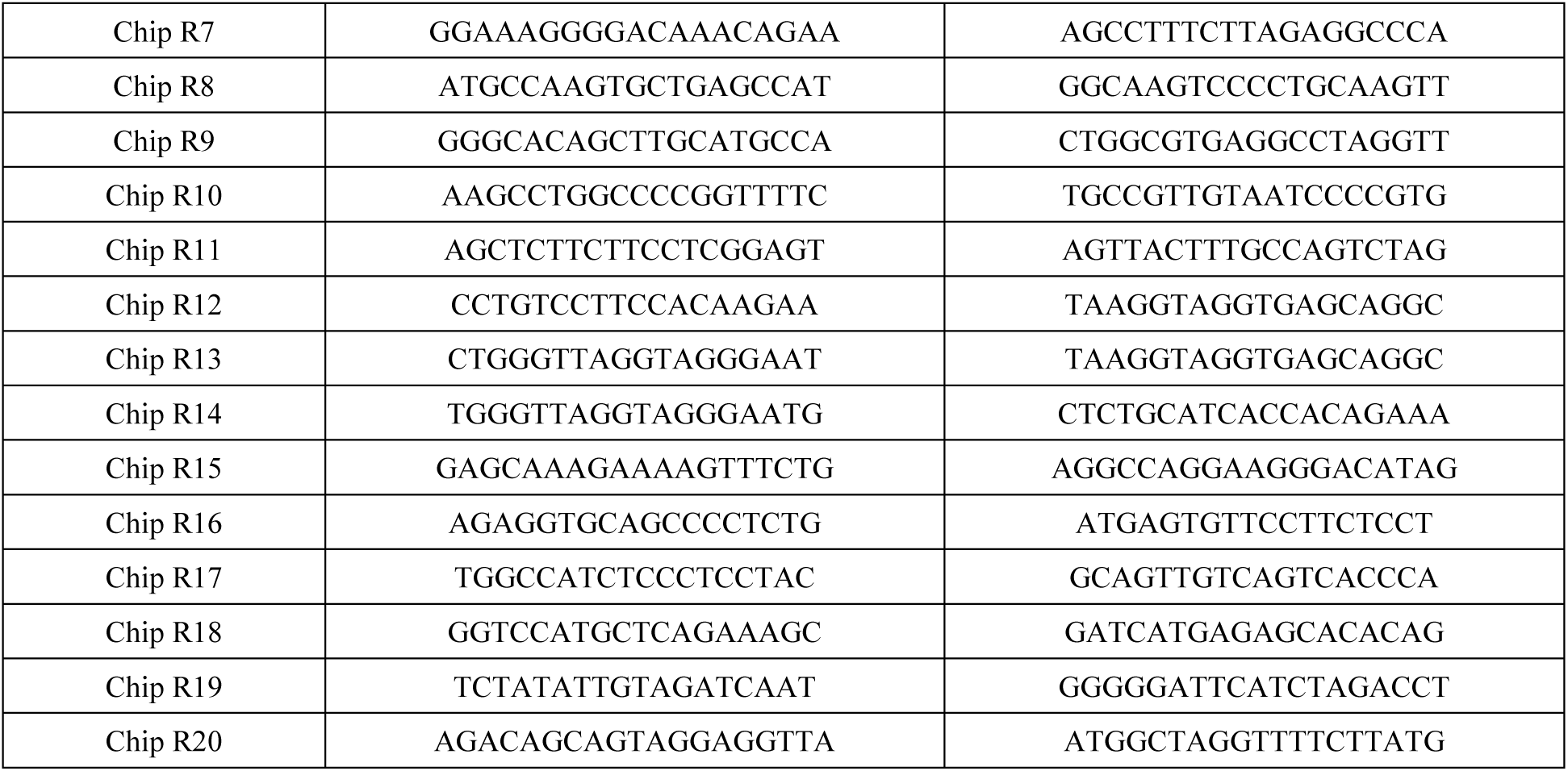
The primers used in Real-time PCR (5’-3’).

## References

1. Ling C, Rönn T. (2019). Epigenetics in Human Obesity and Type 2 Diabetes. Cell Metab 29:1028–44.

2. Klein S, Gastaldelli A, Yki-Järvinen H, Scherer PE. (2022). Why does obesity cause diabetes? Cell Metab 34:11–20.

3. Scherer PE. (2006). Adipose tissue: from lipid storage compartment to endocrine organ. Diabetes 55:1537–45.

4. Kang K, Reilly SM, Karabacak V, Gangl MR, Fitzgerald K, Hatano B, et al. (2008). Adipocyte-derived Th2 cytokines and myeloid PPARdelta regulate macrophage polarization and insulin sensitivity. Cell Metab 7:485–95.

5. Pellegrinelli V, Rodriguez-Cuenca S, Rouault C, Figueroa-Juarez E, Schilbert H, Virtue S, et al. (2022). Dysregulation of macrophage PEPD in obesity determines adipose tissue fibro-inflammation and insulin resistance. Nat Metab 4:476-94.

6. Hägglöf T, Vanz C, Kumagai A, Dudley E, Ortega V, Siller M, et al. (2022). T-bet(+) B cells accumulate in adipose tissue and exacerbate metabolic disorder during obesity. Cell Metab 34:1121–36.e6.

7. Kratz M, Coats BR, Hisert KB, Hagman D, Mutskov V, Peris E, et al. (2014). Metabolic dysfunction drives a mechanistically distinct proinflammatory phenotype in adipose tissue macrophages. Cell Metab 20:614–25.

8. Kohlgruber A, Lynch L. (2015). Adipose tissue inflammation in the pathogenesis of type 2 diabetes. Curr Diab Rep 15:92.

9. Burhans MS, Hagman DK, Kuzma JN, Schmidt KA, Kratz M. (2018). Contribution of Adipose Tissue Inflammation to the Development of Type 2 Diabetes Mellitus. Compr Physiol 9:1–58.

10. Brestoff JR, Wilen CB, Moley JR, Li Y, Zou W, Malvin NP, et al. (2021). Intercellular Mitochondria Transfer to Macrophages Regulates White Adipose Tissue Homeostasis and Is Impaired in Obesity. Cell Metab 33:270–82.e8.

11. Lee BC, Kim MS, Pae M, Yamamoto Y, Eberlé D, Shimada T, et al. (2016). Adipose Natural Killer Cells Regulate Adipose Tissue Macrophages to Promote Insulin Resistance in Obesity. Cell Metab 23:685–98.

12. Weisberg SP, McCann D, Desai M, Rosenbaum M, Leibel RL, Ferrante AW, Jr. (2003). Obesity is associated with macrophage accumulation in adipose tissue. J Clin Invest 112:1796–808.

13. Hotamisligil GS. (2006). Inflammation and metabolic disorders. Nature 444:860–7.

14. Xu H, Barnes GT, Yang Q, Tan G, Yang D, Chou CJ, et al. (2003). Chronic inflammation in fat plays a crucial role in the development of obesity-related insulin resistance. J Clin Invest 112:1821–30.

15. Chawla A, Nguyen KD, Goh YP. (2011). Macrophage-mediated inflammation in metabolic disease. Nat Rev Immunol 11:738–49.

16. Patsouris D, Li PP, Thapar D, Chapman J, Olefsky JM, Neels JG. (2008). Ablation of CD11c-positive cells normalizes insulin sensitivity in obese insulin resistant animals. Cell Metab 8:301–9.

17. Nomiyama T, Perez-Tilve D, Ogawa D, Gizard F, Zhao Y, Heywood EB, et al. (2007). Osteopontin mediates obesity-induced adipose tissue macrophage infiltration and insulin resistance in mice. J Clin Invest 117:2877–88.

18. Arkan MC, Hevener AL, Greten FR, Maeda S, Li ZW, Long JM, et al. (2005). IKK-beta links inflammation to obesity-induced insulin resistance. Nat Med 11:191–8.

19. Patra D, Roy S, Arora L, Kabeer SW, Singh S, Dey U, et al. (2023). miR-210-3p Promotes Obesity-Induced Adipose Tissue Inflammation and Insulin Resistance by Targeting SOCS1-Mediated NF-κB Pathway. Diabetes 72:375–88.

20. Ambros V. (2004). The functions of animal microRNAs. Nature 431:350–5.

21. Dumortier O, Hinault C, Van Obberghen E. (2013). MicroRNAs and metabolism crosstalk in energy homeostasis. Cell Metab 18:312–24.

22. Krützfeldt J, Stoffel M. (2006). MicroRNAs: a new class of regulatory genes affecting metabolism. Cell Metab 4:9–12.

23. Arner P, KulytéA. (2015). MicroRNA regulatory networks in human adipose tissue and obesity. Nat Rev Endocrinol 11:276–88.

24. Thomou T, Mori MA, Dreyfuss JM, Konishi M, Sakaguchi M, Wolfrum C, et al. (2017). Adipose-derived circulating miRNAs regulate gene expression in other tissues. Nature 542:450–5.

25. Agbu P, Carthew RW. (2021). MicroRNA-mediated regulation of glucose and lipid metabolism. Nat Rev Mol Cell Biol 22:425–38.

26. Koh EH, Chernis N, Saha PK, Xiao L, Bader DA, Zhu B, et al. (2018). miR-30a Remodels Subcutaneous Adipose Tissue Inflammation to Improve Insulin Sensitivity in Obesity. Diabetes 67:2541–53.

27. Zhang F, Ma D, Zhao W, Wang D, Liu T, Liu Y, et al. (2020). Obesity-induced overexpression of miR-802 impairs insulin transcription and secretion. Nat Commun 11:1822.

28. Kornfeld JW, Baitzel C, Könner AC, Nicholls HT, Vogt MC, Herrmanns K, et al. (2013). Obesity-induced overexpression of miR-802 impairs glucose metabolism through silencing of Hnf1b. Nature 494:111–5.

29. Ru Y, Kechris KJ, Tabakoff B, Hoffman P, Radcliffe RA, Bowler R, et al. (2014). The multiMiR R package and database: integration of microRNA-target interactions along with their disease and drug associations. Nucleic Acids Res 42:e133.

30. Liao G, Zhang M, Harhaj EW, Sun SC. (2004). Regulation of the NF-kappaB-inducing kinase by tumor necrosis factor receptor-associated factor 3-induced degradation. J Biol Chem 279:26243–50.

31. He L, Grammer AC, Wu X, Lipsky PE. (2004). TRAF3 forms heterotrimers with TRAF2 and modulates its ability to mediate NF-{kappa}B activation. J Biol Chem 279:55855–65.

32. He JQ, Saha SK, Kang JR, Zarnegar B, Cheng G. (2007). Specificity of TRAF3 in its negative regulation of the noncanonical NF-kappa B pathway. J Biol Chem 282:3688–94.

33. Zarnegar B, Yamazaki S, He JQ, Cheng G. (2008). Control of canonical NF-kappaB activation through the NIK-IKK complex pathway. Proc Natl Acad Sci U S A 105:3503–8.

34. Bista P, Zeng W, Ryan S, Bailly V, Browning JL, Lukashev ME. (2010). TRAF3 controls activation of the canonical and alternative NFkappaB by the lymphotoxin beta receptor. J Biol Chem 285:12971–8.

35. Akhter N, Wilson A, Thomas R, Al-Rashed F, Kochumon S, Al-Roub A, et al. (2021). ROS/TNF-α Crosstalk Triggers the Expression of IL-8 and MCP-1 in Human Monocytic THP-1 Cells via the NF-κB and ERK1/2 Mediated Signaling. Int J Mol Sci 22.

36. Cildir G, Low KC, Tergaonkar V. (2016). Noncanonical NF-κB Signaling in Health and Disease. Trends Mol Med 22:414–29.

37. Shimano H, Sato R. (2017). SREBP-regulated lipid metabolism: convergent physiology - divergent pathophysiology. Nat Rev Endocrinol 13:710–30.

38. Lumeng CN, Bodzin JL, Saltiel AR. (2007). Obesity induces a phenotypic switch in adipose tissue macrophage polarization. J Clin Invest 117:175–84.

39. Prieur X, Mok CY, Velagapudi VR, Núñez V, Fuentes L, Montaner D, et al. (2011). Differential lipid partitioning between adipocytes and tissue macrophages modulates macrophage lipotoxicity and M2/M1 polarization in obese mice. Diabetes 60:797–809.

40. Ge W, Goga A, He Y, Silva PN, Hirt CK, Herrmanns K, et al. (2022). miR-802 Suppresses Acinar-to-Ductal Reprogramming During Early Pancreatitis and Pancreatic Carcinogenesis. Gastroenterology 162:269–84.

41. Goga A, Yagabasan B, Herrmanns K, Godbersen S, Silva PN, Denzler R, et al. (2021). miR-802 regulates Paneth cell function and enterocyte differentiation in the mouse small intestine. Nat Commun 12:3339.

42. Gao T, Zou M, Shen T, Duan S. (2021). Dysfunction of miR-802 in tumors. J Clin Lab Anal 35:e23989.

43. Seok S, Sun H, Kim YC, Kemper B, Kemper JK. (2021). Defective FXR-SHP Regulation in Obesity Aberrantly Increases miR-802 Expression, Promoting Insulin Resistance and Fatty Liver. Diabetes 70:733–44.

44. Ni Y, Xu Z, Li C, Zhu Y, Liu R, Zhang F, et al. (2021). Therapeutic inhibition of miR-802 protects against obesity through AMPK-mediated regulation of hepatic lipid metabolism. Theranostics 11:1079–99.

45. Häcker H, Tseng PH, Karin M. (2011). Expanding TRAF function: TRAF3 as a tri-faced immune regulator. Nat Rev Immunol 11:457–68.

46. Lumeng CN, Deyoung SM, Bodzin JL, Saltiel AR. (2007). Increased inflammatory properties of adipose tissue macrophages recruited during diet-induced obesity. Diabetes 56:16–23.

47. Flaherty SE, 3rd, Grijalva A, Xu X, Ables E, Nomani A, Ferrante AW, Jr. (2019). A lipase-independent pathway of lipid release and immune modulation by adipocytes. Science 363:989-93.

48. Batista-Gonzalez A, Vidal R, Criollo A, Carreño LJ. (2019). New Insights on the Role of Lipid Metabolism in the Metabolic Reprogramming of Macrophages. Front Immunol 10:2993.

49. Im SS, Yousef L, Blaschitz C, Liu JZ, Edwards RA, Young SG, et al. (2011). Linking lipid metabolism to the innate immune response in macrophages through sterol regulatory element binding protein-1a. Cell Metab 13:540–9.

50. Fei X, Huang J, Li F, Wang Y, Shao Z, Dong L, et al. (2023). The Scap-SREBP1-S1P/S2P lipogenesis signal orchestrates the homeostasis and spatiotemporal activation of NF-κB. Cell Rep 42:112586.

51. Virtue S, Vidal-Puig A. (2021). GTTs and ITTs in mice: simple tests, complex answers. Nat Metab 3:883-6.

52. Gordon DM, Neifer KL, Hamoud AA, Hawk CF, Nestor-Kalinoski AL, Miruzzi SA, et al. (2020). Bilirubin remodels murine white adipose tissue by reshaping mitochondrial activity and the coregulator profile of peroxisome proliferator-activated receptor α. J Biol Chem 295:9804–22.

